# Genome-wide imaging screen uncovers molecular determinants of arsenite-induced protein aggregation and toxicity and an important role for transcriptional and translational control

**DOI:** 10.1101/2020.12.16.423032

**Authors:** Stefanie Andersson, Antonia Romero, Joana Isabel Rodrigues, Sansan Hua, Xinxin Hao, Therese Jacobson, Vivien Karl, Nathalie Becker, Arghavan Ashouri, Thomas Nyström, Beidong Liu, Markus J. Tamás

**Author notes:** Corresponding author: Markus J. Tamás, Department of Chemistry and Molecular Biology, University of Gothenburg, S-405 30 Göteborg, Sweden, Tel: -46-31-786 2548. These authors contributed equally to the study (Stefanie Andersson, Antonia Romero).

## Abstract

Exposure to toxic metals and metalloids such as cadmium and arsenic results in widespread misfolding and aggregation of cellular proteins. How these protein aggregates are formed *in vivo*, the mechanisms by which they affect cells, and how cells prevent their accumulation during environmental stress is not fully understood. To find components involved in these processes, we performed a genome-wide imaging screen and identified yeast deletion mutants with either enhanced or reduced protein aggregation levels during arsenite exposure. Mutants with reduced aggregation levels were enriched for functions related to protein biosynthesis and transcription, whilst functions related to cellular signalling, metabolism, and protein folding and degradation were overrepresented among mutants with enhanced aggregation levels. On a genome-wide scale, protein aggregation correlated with arsenite resistance and sensitivity, indicating that many of the identified factors are crucial to safeguard protein homeostasis (proteostasis) and to protect against arsenite toxicity. Dedicated follow-up experiments indicated that intracellular arsenic is a direct cause of protein aggregation and that accurate transcriptional and translational control are crucial for proteostasis during arsenite stress. Specifically, we provide evidence that global transcription affects protein aggregation levels, that loss of transcriptional control impacts proteostasis through distinct mechanisms, and that translational repression is central to control protein aggregation and cell viability. Some of the identified factors are associated with pathological conditions suggesting that arsenite-induced protein aggregation may impact disease processes. The broad network of cellular systems that impinge on proteostasis during arsenic stress provides a valuable resource and a framework for further elucidation of the mechanistic details of metalloid toxicity and pathogenesis.

**AUTHOR SUMMARY:** Human exposure to poisonous metals is increasing in many parts of the world and chronic exposure is associated with certain protein folding-associated disorders such as Alzheimer’s disease and Parkinson’s disease. While the toxicity of many metals is undisputed, their molecular modes of action have remained unclear. Recent studies revealed that toxic metals such as arsenic and cadmium profoundly affect the correct folding of proteins, resulting in the accumulation of toxic protein aggregates. In this study, we used high-content microscopy to identify a broad network of cellular systems that impinge on protein homeostasis and cell viability during arsenite stress. Follow-up experiments highlight the importance of accurate transcriptional and translational control for mitigating arsenite-induced protein aggregation and toxicity. Some of the identified factors are associated with pathological conditions suggesting that arsenite-induced protein aggregation may impact disease processes. The broad network of cellular systems that impinge on proteostasis during arsenic stress provides a valuable resource and a framework for further elucidation of the mechanistic details of metal toxicity and pathogenesis.

## INTRODUCTION

The processes that control protein synthesis, folding, localization, abundance and degradation are crucial for the proper functioning of cells and organisms. The ability of the cell to maintain a functional proteome (protein homeostasis or proteostasis) declines during ageing which may lead to the accumulation of damaged, misfolded and aggregated proteins. The cellular proteome is also threatened by environmental stress conditions that promote rapid and extensive protein misfolding and aggregation. Excessive protein misfolding and aggregation can cause cellular or organismal damage, as exemplified by the many pathological conditions that are associated with defective proteostasis, including neurodegenerative and age-related disorders such as Alzheimer’s disease (AD) and Parkinson’s disease (PD) [1–4]. How different aggregated protein species cause or contribute to toxicity or pathology is poorly understood.

Cells use a battery of protein quality control (PQC) mechanisms to ensure proteostasis. Molecular chaperones assist the folding of proteins into their functional conformation or help misfolded proteins to regain their native structures. Molecular chaperones are also integral parts of protein degradation systems, such as the proteasome, the autophagy pathway, and the lysosome/vacuole, that clear cells from aberrant protein conformers [1–4]. Misfolded and aggregated proteins may be directed to specific subcellular deposition sites, as a means to reduce their toxicity [5, 6]. Cells may also use the controlled formation of protein aggregates for various physiological purposes [7, 8] such as storage of peptide and protein hormones [9], regulation of cell cycle restart after stress [10] and microbial adhesion, biofilm formation and host invasion [8, 11]. The potential toxicity or benefit of protein aggregates highlights the importance of proteostasis during physiological conditions as well as during ageing, pathological conditions and stress.

Human exposure to poisonous metals is increasing in many parts of the world and chronic exposure is associated with certain protein folding-associated diseases including AD and PD [12–15]. While the toxicity of many metals is undisputed, their molecular modes of action have remained unclear. Recent *in vitro* and *in vivo* studies revealed that toxic metals profoundly affect proteostasis [16–22]. Arsenic, in form of trivalent arsenite [As(III)] [18, 23], and cadmium [19] were shown to cause widespread protein aggregation in living yeast cells primarily by targeting nascent proteins. *In vitro* and *in vivo* data suggested that As(III)- and cadmium-aggregated protein species may form seeds that increase the misfolding and aggregation of other susceptible proteins. These studies also suggested that misfolding and aggregation of nascent proteins represents an important component of arsenite’s and cadmium’s mode of toxic action [18, 19]. Toxicity of chromium, in form of Cr(VI), is partly a result of enhanced mRNA mistranslation leading to the accumulation of misfolded and aggregated proteins [21]. Despite of these examples, our current knowledge of the molecular basis of metal stress-induced protein aggregation in living cells and how cells regulate the PQC systems to protect against toxic aggregates is incomplete. In this study, we used high-content microscopy to identify a broad network of cellular systems that impinge on proteostasis and cell viability during arsenite stress. Follow-up experiments highlight the importance of accurate transcriptional and translational control for mitigating arsenite-induced protein aggregation and toxicity.

## RESULTS

### Genome-wide imaging screen uncovers yeast deletion mutants with either enhanced or reduced levels of protein aggregation during As(III) exposure

To identify factors that impinge on proteostasis during As(III) exposure, we performed a high-content imaging screen in the budding yeast *Saccharomyces cerevisiae*. For this, we first incorporated a GFP (green fluorescence protein) tagged version of Hsp104, a molecular chaperone that binds to and disassembles protein aggregates [24], into a genome-wide collection of viable deletion mutants using a synthetic genetic array (SGA) approach (Fig. 1A). Hsp104-GFP can be used as a visual marker during As(III) exposure as it redistributes from a diffuse cytosolic localization to specific foci that represent protein aggregates [18]. We cultivated and exposed this strain collection to As(III) using high-throughput robotic handling in a microtiter plate format. The percentage of cells containing Hsp104-GFP foci/protein aggregates was systematically scored using automated image analysis and quantitated as the deviation from the aggregation level in wild type cells (see Materials and Methods). To allow identification of mutants that accumulate either more or fewer protein aggregates than the wild type control, we chose an As(III) concentration (0.25 mM) and a time point (2 h) at which ~40% of the wild type cells contained Hsp104-GFP foci. We counted cells that had 1-2 aggregates/cell and those with ≥3 aggregates/cell, and the obtained values for each mutant were then compared to those for wild type cells. Mutants that deviated significantly (*p*<0.05) in aggregate levels (fraction of cells with aggregates) from the wild type were selected for further analysis (Supplemental data Table S1). In this way, we found 202 mutants that accumulated more aggregates (*i.e*. showing enhanced aggregation) and 198 mutants with fewer aggregates than the wild type (reduced aggregation) (Fig. 1A).

**Figure 1.**
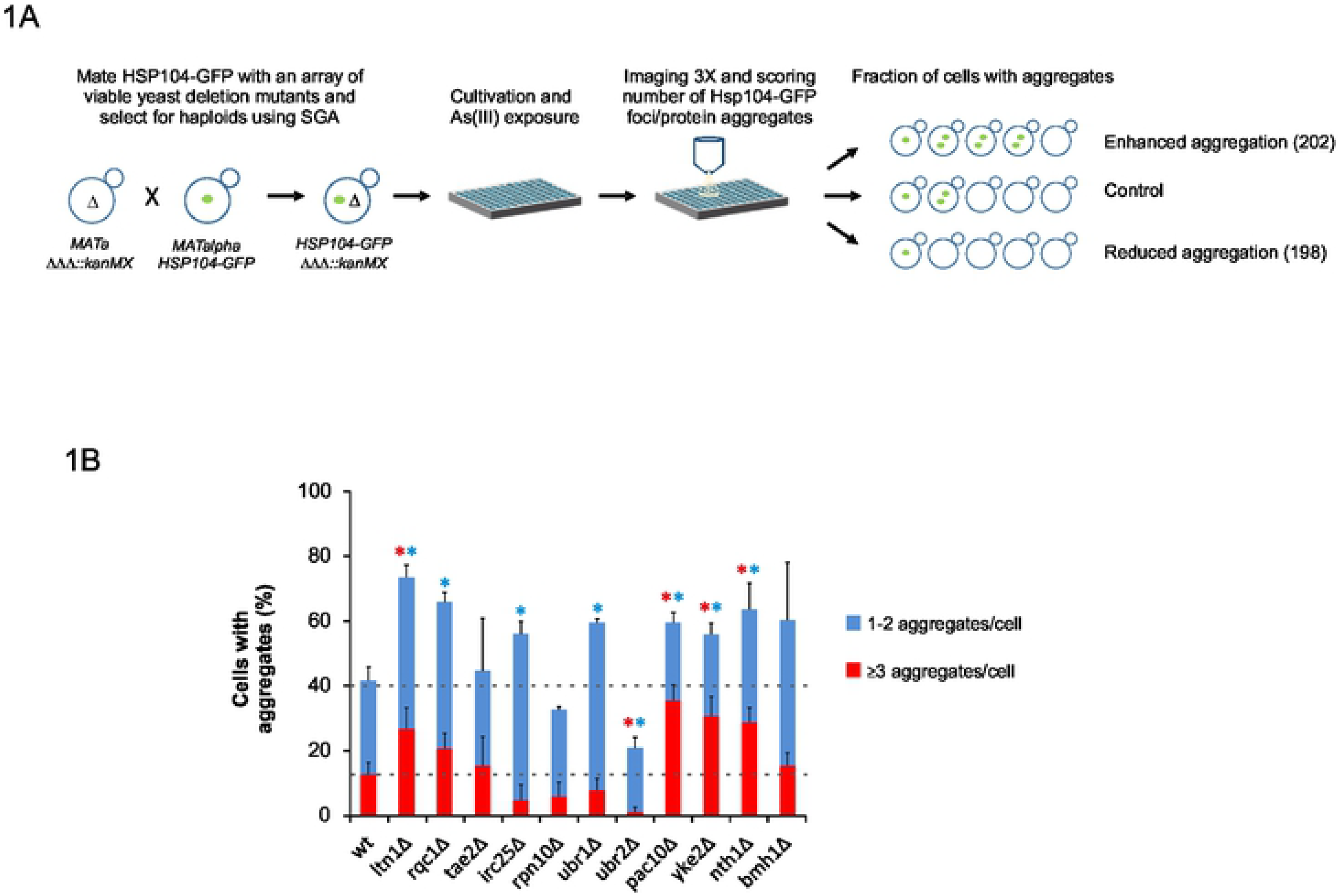
Systematic high-content screen identifies molecular determinants of As(III)-induced protein aggregation. **1A)** Schematic representation of the high-content imaging screen workflow. Hits from the screen were divided into two categories: mutants that accumulated more aggregates than the wild type (enhanced aggregation) and mutants with fewer aggregates than the wild type (reduced aggregation). **1B)** Validation and quantification of protein aggregation in selected mutants. Hsp104-GFP distribution was scored in wild type and mutant cells after 3 h of As(III) exposure (0.5 mM). The fractions of cells containing 1-2 aggregates/cell and ≥3 aggregates/cell were determined by visual inspection of ~200 cells per strain. Error bars represent standard deviations (S.D.) from two (mutants) and six (wild type) independent biological replicates. The error bars on the top concern the total fraction of cells with aggregates, whilst those on the red bars concern the fraction of cells with ≥3 aggregates/cell. Statistical analyses were performed by Student’s t-test and significant differences are indicated by a star (p<0.01); blue star: total fraction of cells with aggregates; red star: ≥3 aggregates/cell.

The large number of hits suggests that yeast devotes a substantial fraction (~400 hits represent ~8% of non-essential *S. cerevisiae* genes) of its genome into PQC during As(III) stress. Among the hits, several were previously known or expected to affect proteostasis (see below), validating the screening approach. To directly test the accuracy of the screen, we manually re-created 11 strains by crossing individual mutants selected from our hit list with an Hsp104-GFP strain, exposed the resulting strains to 0.5 mM As(III) (a higher As(III) concentration is needed to obtain similar aggregation levels in flasks versus microtiter plates), and scored their respective protein aggregation levels. 8 out of the 11 strains deviated significantly (*p*<0.01) from the protein aggregation levels in wild type cells at 3 h of As(III) exposure (Fig. 1B). Using other assays, we confirmed another 8 mutants out of 12 tested (Figs 6, 7), suggesting an overall accuracy of at least 70% (true positives). Hence, the identified hits likely represent a comprehensive catalogue of factors regulating proteostasis during As(III) stress. The gene products that are directly involved in proteostasis remain to be distinguished from those with an indirect effect.

**Figure 6.**
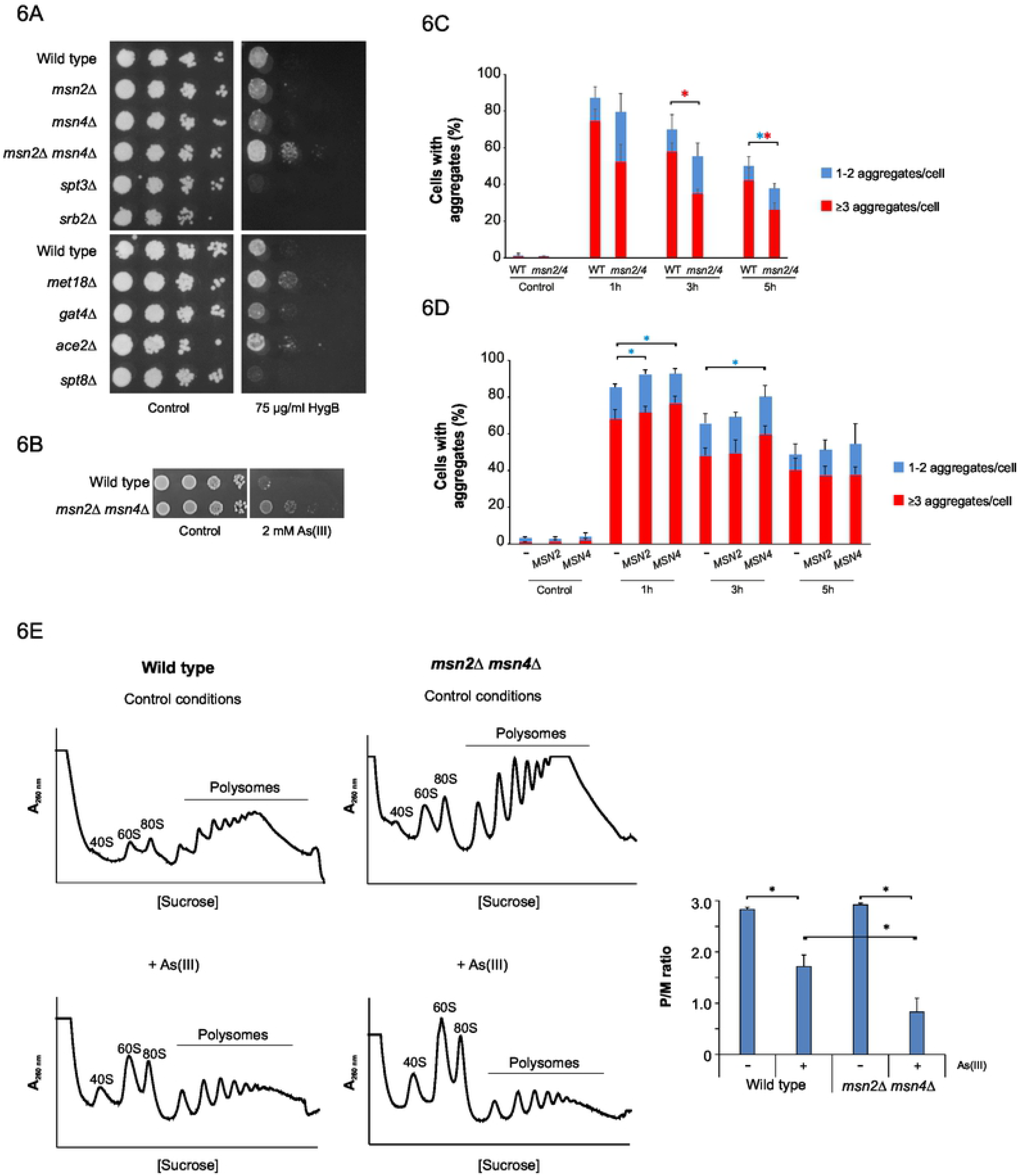
Loss of positive regulators of transcription leads to reduced protein aggregation. **6A)** 10-fold serial dilutions of the indicated strains were plated onto YPD agar plates with or without HygB. Growth was monitored after 2-3 days at 30°C. **6B)** 10-fold serial dilutions of the indicated strains were plated onto agar plates with or without As(III). Growth was monitored after 2-3 days at 30°C. **6C)** Quantification of protein aggregation. Sis1-GFP distribution was scored in wild type and *msn2Δ msn4Δ* cells by fluorescence microscopy before and after exposure to 0.5 mM As(III). The fractions of cells containing 1-2 aggregates/cell and ≥3aggregates/cell were determined by visual inspection of 102-131 cells per condition and time-point. Error bars represent standard deviations (s.d.) from five independent biological replicates (n=5). The error bars on the top concern the total fraction of cells with aggregates, whilst those on the red bars concern the fraction of cells with ≥3 aggregates/cell. Statistical analyses were performed by Student’s t-test and significant differences are indicated by star (*p*<0.05). Blue star: total fraction of cells with aggregates; red star: ≥3 aggregates/cell. **6D)** Quantification of protein aggregation. Hsp104-GFP distribution was scored as described above (Fig. 6C) in wild type cells transformed with an empty plasmid (-) or with centromeric plasmids harbouring *MSN2* or *MSN4*. **6E)** Lack of stress-responsive activators Msn2 and Msn4 results in efficient translation inhibition in response to As(III). Polysome profiles were obtained from wild type and *msn2Δ msn4Δ* strains cultivated in synthetic complete medium for at least 3 h to reach exponential phase, and then cells were further maintained in or treated with 0.75 mM As(III) for 1 h. The A_260nm_ profiles after gradient fractionation are shown and the ribosomal subunits (40S and 60S), monosomes (80S) and polysomes are indicated. Three biologically independent replicates were performed in each case (n=3). A representative profile is shown. Right panel: the average polysome/monosome ratio (P/M) is represented with its corresponding standard deviation. Star indicates statistically significant difference (*p*<0.05).

**Figure 7.**
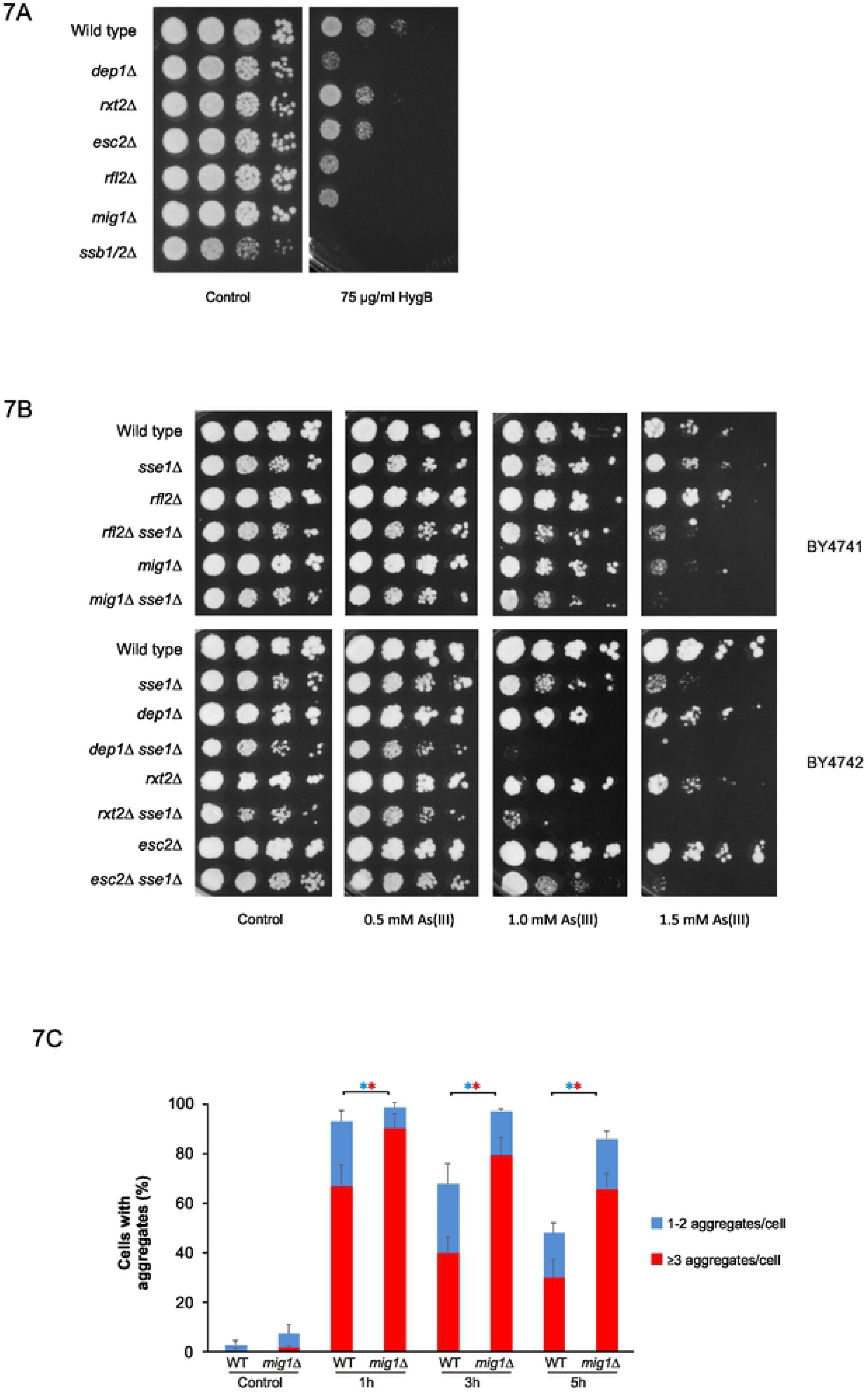
Loss of negative regulators of transcription leads to enhanced protein aggregation. **7A)** 10-fold serial dilutions of the indicated strains were plated onto YPD agar plates with or without HygB. Growth was monitored after 2-3 days at 30°C. **7B)** Loss of Sse1 exacerbates As(III) sensitivity of mutants that lack negative regulators of transcription. 10-fold serial dilutions of the indicated cells were plated onto YPD agar plates with or without As(III). Growth was monitored after 2-3 days at 30°C. **7C)** Quantification of protein aggregation. Sis1-GFP distribution was scored in wild type and *mig1Δ* cells by fluorescence microscopy before and after exposure to 0.5 mM As(III). The fractions of cells containing 1-2 aggregates/cell and ≥3aggregates/cell were determined by visual inspection of 111-161 cells per condition and time-point. Error bars represent standard deviations (S.D.) from three independent biological replicates (n=3). The error bars on the top concern the total fraction of cells with aggregates, whilst those on the red bars concern the fraction of cells with ≥3 aggregates/cell. Statistical analyses were performed by Student’s t-test and significant differences are indicated by star (*p*<0.05). Blue star: total fraction of cells with aggregates; red star: ≥3 aggregates/cell.

### Loss of functions related to transcription and translation leads to reduced protein aggregation levels

We next investigated whether specific categories of protein functions were over-represented in our data-sets. The set of mutants that accumulated fewer protein aggregates than the wild type was primarily enriched (according to FunCat, Munich Information Center for Protein Sequences (MIPS) [25]) for genes with functions in protein biosynthesis, protein and RNA binding, and for cytosolic proteins (Fig. 2A; Table S1). To identify biologically meaningful connections between these proteins, we constructed an interaction network using the STRING (Search Tool for Retrieval of Interacting Genes/Proteins) database [26, 27]. A large proportion of the hits are part of highly connected networks (protein-protein interaction (PPI) enrichment *p*-value <10^−16^) involved in cytoplasmic translation (*e.g.* ribosomal proteins), rRNA modification and processing, ribosome assembly, as well as several members of the elongator (ELP) complex that is primarily involved in tRNA modifications [28] (Supplemental Fig. S1; Table S1). We next visualized the functional diversity of our gene-list by mapping each hit onto the global yeast genetic interaction network using CellMap [29, 30]. This analysis showed that the hits are clustered in distinct functions related to chromatin, transcription, ribosome biogenesis, rRNA processing, mRNA processing, tRNA wobble modification as well as glycosylation, protein folding and cell wall (Fig 2C). Together, the hits support earlier observations that processes related to translation impinge on proteostasis during As(III) stress [18, 23, 31]. Our analysis also identified protein modules that were previously not linked to protein aggregation including proteins with functions in mRNA modification, transcription, and chromatin organization (Fig. S1; Table S1). Thus, transcription-related processes may contribute to aggregate management during metalloid exposure (see further).

**Figure 2.**
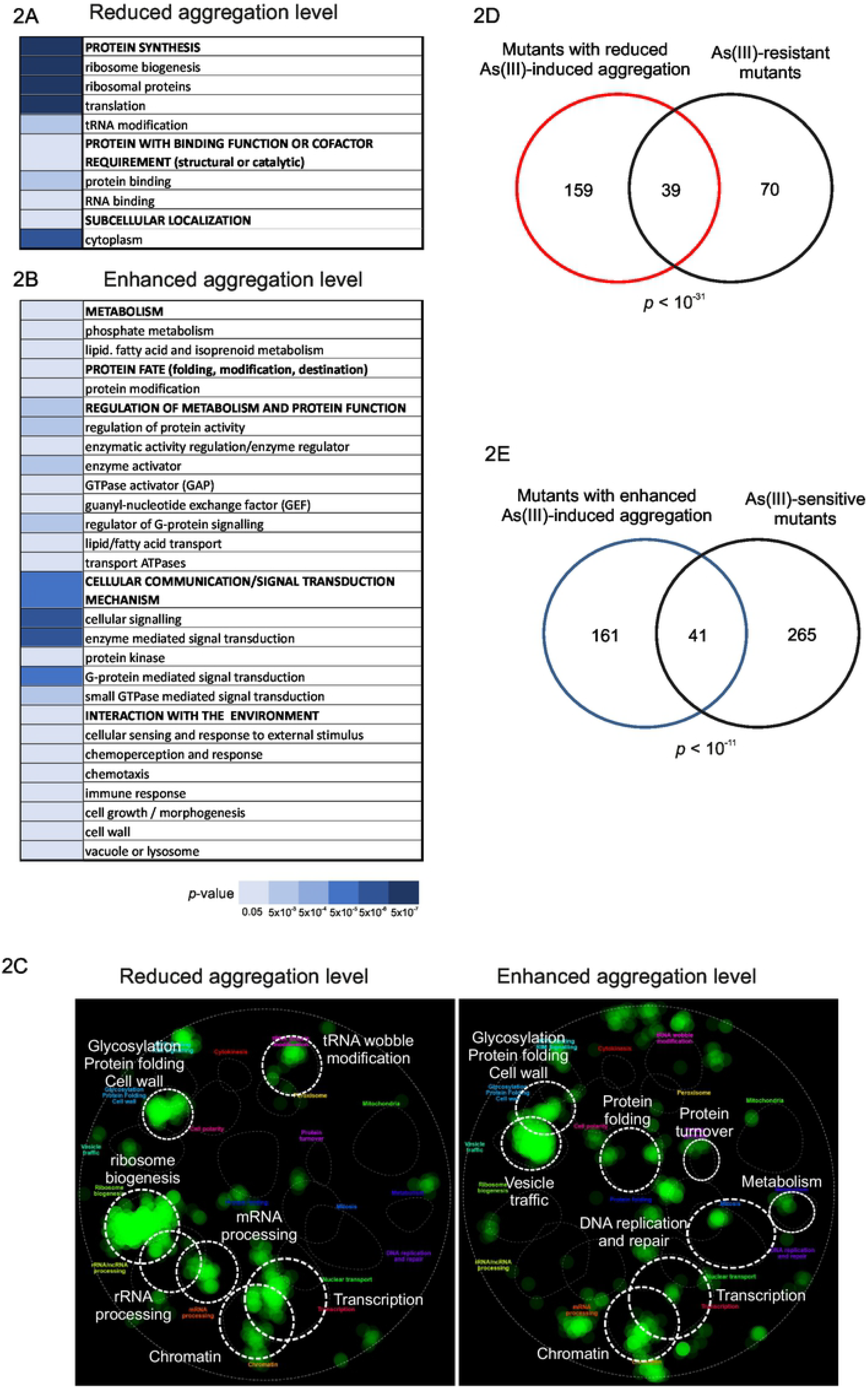
Characterization of the hits. Functional categories of hits (according to MIPS) that were significantly enriched (FDR<0.005) among mutants with reduced (**2A**) and enhanced (**2B**) aggregation. (**2C**) The identified hits belong to distinct functions. Each hit was mapped onto the global yeast genetic interaction network and visualized using CellMap [29, 30]. Correlation between protein aggregation and growth in the presence of As(III). The Venn diagrams show the overlap between: **(2D)** mutants with reduced level of As(III)-induced aggregation and mutants that show As(III)-resistance [37]; and between **(2E)** mutants with enhanced level of As(III)-induced aggregation and mutants that show As(III)-sensitivity [38]. Significance of the overlaps was calculated by the hyper-geometric test and the corresponding *p*-values are indicated.

### Mutants with enhanced levels of protein aggregation are enriched in a range of cellular functions

The set of mutants with enhanced levels of protein aggregation was enriched (according to MIPS) in a range of biological functions including metabolism (primarily phosphate and lipid metabolism), protein fate (folding, modification, destination), regulation of metabolism and protein function, cellular communication and signalling, as well as interaction with the environment (Fig. 2B; Table S1). The hits are part of highly connected networks (PPI enrichment *p*-value 10^−14^) associated with protein folding and degradation, signalling, transcription, and lipid/fatty acid metabolism (Fig. S2). Mapping these hits onto the global yeast genetic interaction network (CellMap) identified clusters of functions related to chromatin, transcription, metabolism, DNA replication and repair, protein folding and turnover, vesicle traffic, as well as glycosylation, protein folding and cell wall (Fig. 2C). Protein folding-associated modules included members of the prefoldin complex (Pac10, Yke2, Gim4) involved in actin and tubulin folding as well as in transcription elongation [32], and members of the GET complex (Get1, Get2, Get3, Get4) that promotes insertion of tail-anchored proteins into the ER membrane (Fig. S2). Get3 also functions as a holdase chaperone under oxidative stress conditions [33, 34]. The presence of protein folding-related genes in the enhanced aggregation set supports previous findings that As(III) interferes with protein folding processes *in vivo* [18, 23]. Protein turnover-associated hits included members of the ribosomal quality control (RQC) complex (Ltn1, Rqc1) that mediate the recognition and ubiquitylation of ribosome-associated aberrant nascent polypeptides [35], several components of the ubiquitin-proteasome pathway, as well as proteins involved in intracellular trafficking including a number of autophagy-related proteins (Fig. S2). Thus, clearance of As(III)-induced protein aggregates may involve both the ubiquitin-proteasome and the autophagy pathways, in line with earlier reports [18, 31]. A large number of genes previously not linked to protein aggregation were also identified. For example, network analyses pin-pointed protein modules involved in signalling (Snf1 and glucose/cAMP signalling pathways, the Hog1 mitogen-activated protein (MAP) kinase pathway, and G-protein-mediated signalling), and transcription including members of the Rpd3 histone deacetylase complex (Fig. S2). Hence, signal transduction and transcriptional regulatory pathways may impinge on PQC during stress. We also identified genes known to have direct roles in arsenic detoxification, as well as a number of genes with functions in lipid/fatty acid metabolism (Table S1; Fig. S2). A few genes with translation-related functions were found among mutants with enhanced aggregation levels whilst the majority of protein synthesis-related genes were present in the set with decreased aggregation (Fig. S1). It is possible that this sub-set of proteins has specific functions during translation and their absence might result in imperfect or increased translation, which in turn may lead to increased protein misfolding. Accordingly, cells lacking *ASC1*, which is a negative regulator of translation [36], exhibited enhanced aggregation during As(III) stress (Table S1; Fig. S2).

### Correlation between protein aggregation levels and As(III) sensitivity

Protein aggregates may be toxic or beneficial. To address, in an unbiased manner, whether the aggregates formed during As(III) stress contribute to toxicity, we compared the sets of mutants with reduced and enhanced aggregation to sets of mutants identified in genome-wide As(III) sensitivity and resistance screens. We first compared the overlap between the 198 mutants with reduced aggregation levels to a set of 109 mutants that were previously shown to be As(III) resistant [37]. Importantly, we found a significant overlap between the two gene-sets (39 genes, *p*<10^−31^) (Fig. 2D). Likewise, we observed a significant overlap (41 genes, *p*<10^−11^) between the 202 mutants with enhanced aggregation levels and 306 mutants that were previously shown to be As(III) sensitive [38] (Fig. 2E). In contrast, the overlaps were small between the enhanced aggregation and As(III) resistance sets (7 genes, *p*=0.04) as well as between the reduced aggregation and As(III) sensitivity sets (5 genes, *p*=0.01). Thus, on a genome-wide scale, enhanced protein aggregation correlated with As(III) sensitivity, whilst As(III) resistance correlated with reduced aggregation levels. These findings support the notion that protein misfolding and aggregation contributes to the toxicity of this metalloid.

### Intracellular arsenic is a direct cause of protein aggregation

Genes related to arsenic transport and detoxification were present in both the enhanced and reduced aggregation sets (Figs. 3A, S1, S2; Table S1). Mutants lacking Fps1, the aquaglyceroporin that mediates As(III) entry into cells [39], as well as mutants lacking positive regulators of Fps1 (Rgc1, Rgc2 [40], Slt2 [41]) exhibited diminished aggregation levels (Fig. 3A; Table S1). Conversely, cells lacking the As(III) export protein Acr3 [42, 43] or Yap8, the transcription factor that regulates *ACR3* expression [44, 45], had more aggregates than the wild type. Cells lacking Ycf1, the transporter that catalyses As(III) sequestration into vacuoles [39, 43], or Yap1, the transcription factor that regulates *YCF1* expression [44], had more aggregates than the wild type. Cells lacking Ybp1, a positive regulator of Yap1 [46], exhibited enhanced aggregation levels. Mutants lacking the MAP kinase Hog1, a negative regulator of Fps1 activity [47], as well as its upstream components (Pbs2, Ssk1, Ssk2), accumulated more aggregates whilst cells without Ptp3, a negative regulator of Hog1 activity, had less aggregates compared to wild type (Fig. 3A; Table S1). Deletion of some of these genes were previously shown to affect intracellular arsenic levels: *fps1Δ* cells accumulated less arsenic [39] whilst *acr3Δ* and *hog1Δ* accumulated more [47, 48] than wild type cells. These results raised the possibility that protein aggregation may be correlated with intracellular arsenic levels. Uptake measurements confirmed that *acr3Δ* and *hog1Δ* cells accumulate more whilst *fps1Δ* cells accumulate less intracellular arsenic than wild type cells (Fig. 3B). In addition, *yap1Δ*, *yap8Δ* and *ycf1Δ* cells accumulated more arsenic than the wild type, whereas cells lacking the Fps1 regulators Rgc1 and Rgc2 (*rgc1/2Δ*) accumulated less arsenic. Less arsenic was taken up by cells lacking the two phosphatases Ptp2 and Ptp3 (*ptp2/3Δ*) that act as negative regulators of Hog1 (Fig. 3B). Thus, increased intracellular arsenic accumulation results in enhanced protein aggregation levels whilst reduced intracellular arsenic mitigates protein aggregation, suggesting that intracellular arsenic is a direct cause of protein aggregation.

**Figure 3.**
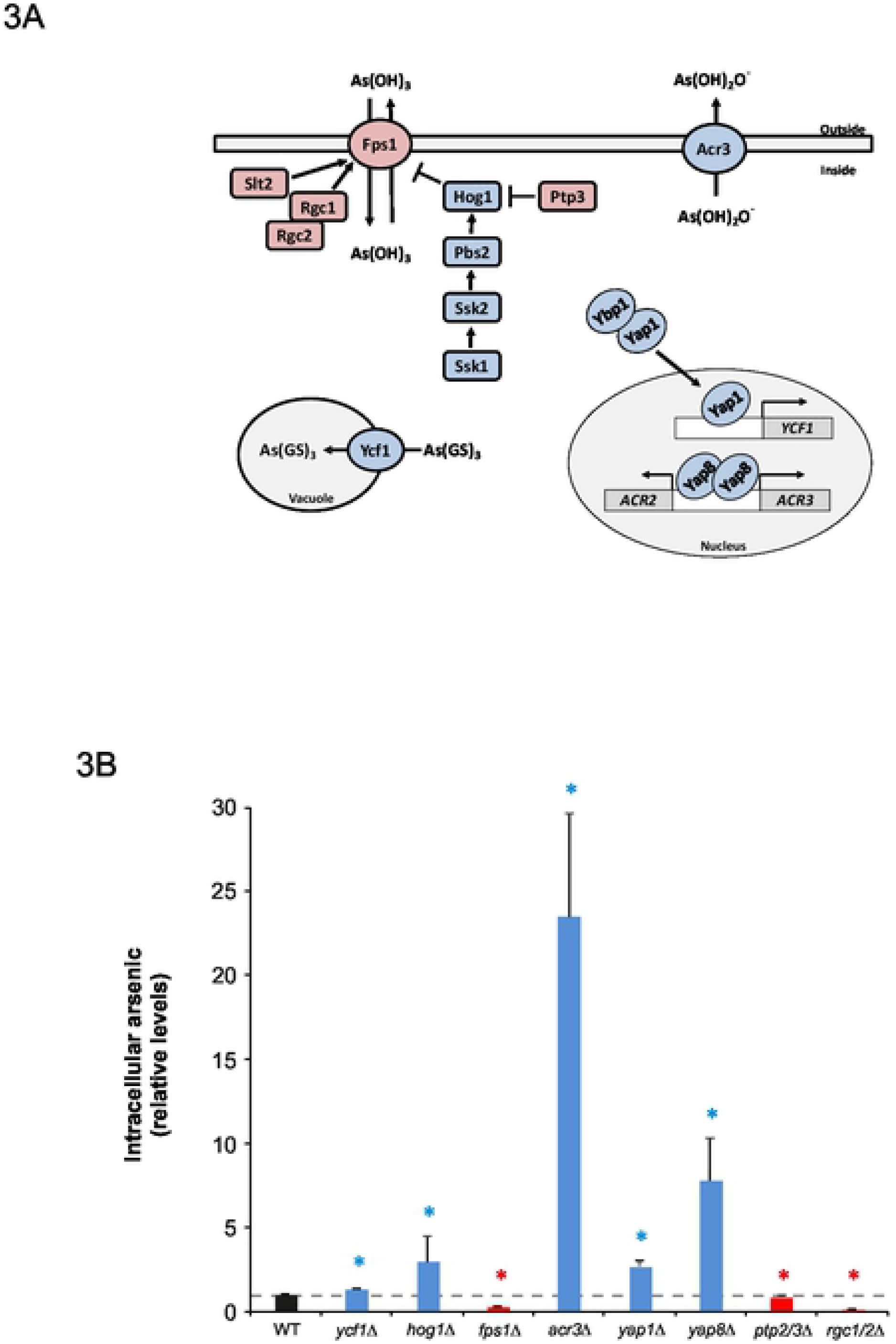
Correlation intracellular arsenic vs. protein aggregation. **3A)** Cartoon showing proteins that regulate arsenic transport and intracellular arsenic levels. Loss of proteins coloured red reduces protein aggregation whilst loss of blue-coloured proteins enhances protein aggregation levels during As(III) exposure. **3B)** Intracellular arsenic. Cells were exposed to As(III) for 1 h, and intracellular arsenic levels were determined as described in Materials and Methods. The data shown represents the average of at least three independent biological replicates and the error bars represent the S.D. Red bars represent mutants with less intracellular arsenic whilst blue bars represent mutants with more intracellular arsenic compared to wild type cells. Relative values are shown to enable comparison between strain backgrounds. Statistical analyses were performed by Student’s t-test and significant differences are indicated by * (*p*<0.05).

### As(III) does not induce transcription errors

Several transcription-related genes were found among mutants with reduced as well as enhanced aggregation levels (Fig. 2; Table S1). Most of these genes have previously not been associated with protein aggregation and the impact of transcriptional control on proteostasis is largely unexplored. One way As(III) could trigger protein aggregation is by causing errors during transcription. Transcription errors have been shown to induce proteotoxic stress and to shorten the lifespan of yeast [49]. To test whether As(III) causes an increased frequency of transcriptional errors, we used an established genetic, assay that detects transcription errors by a Cre-dependent rearrangement of a *HIS3*-based reporter gene [50]. Patches of cells were grown on rich YPD medium and then replica plated onto synthetic medium lacking histidine (His^−^), either in the presence or absence of As(III). On His^−^ plates, His^+^ cells that arise due to transcription errors can form colonies and grow within the patch. Under non-stress conditions, few wild type cells grew on medium lacking histidine (Fig. 4A) indicating a low transcriptional error rate. In the presence of As(III), only few colonies grew on His^−^ plates indicating that As(III) does not promote transcription errors. To substantiate this finding, we repeated the experiment with cells lacking Rpb9 (r*pb9Δ*) since this mutant displays reduced transcription fidelity and elevated error frequency [50]. Accordingly, many colonies of *rpb9Δ* grew on His^−^ plates under non-stress conditions (Fig. 4A). Like for the wild type, As(III) did not affect the number of r*pb9Δ* colonies on His^−^ medium compared to non-stress conditions. In addition to these qualitative assays, we performed quantitative assays where cells were grown in liquid YPD medium and then plated on selective His^−^ medium with or without As(III), as well as on YPD plates to quantify the mean frequency of His^+^ cells relative to the total number of viable cells. Again, there was no significant increase in the frequency of His^+^ cells during As(III) exposure compared to untreated cells, neither in wild type nor *rpb9Δ* cells (Fig. 4B). Thus, As(III) does not induce transcription errors.

**Figure 4.**
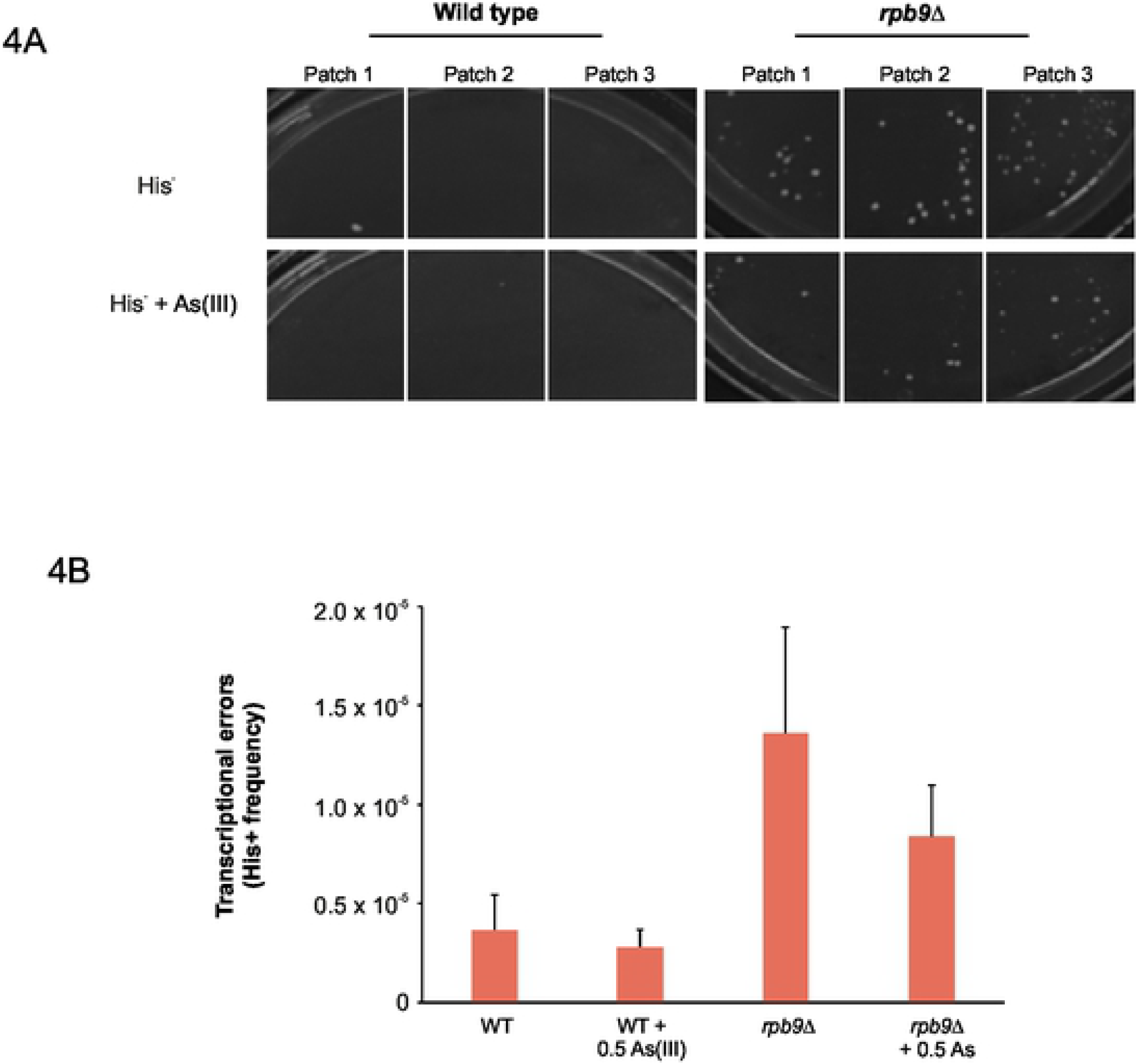
As(III) does not induce transcription errors. **4A)** P_*GAL1*_-*cre*-*Y324C* strains (wild type and *rpb9Δ*) were grown on YPD agar plates overnight at 30°C and replica plated onto synthetic media lacking histidine (His^−^) with or without 0.5 mM As(III). Transcription error results in growth of colonies within a patch on His^−^ plates. 3 representative patches are shown for each strain. Pictures were taken after 3 days of growth at 30°C. **4B)** Mean frequency of His^+^ cells relative to total viable cells in P_*GAL1*_-*cre*-*Y324C* strains. Cultures were grown overnight in 1 ml of YPD at 30°C and plated on SC-His^−^ and YPD agar plates in 10-fold dilutions. Individual colonies were scored after 3 days of growth at 30°C. Results shown are the mean of three independent biological replicates and error bars represent standard error (S.E.).

### Global transcription affects protein aggregation levels during As(III) stress

A subset of the hits with reduced aggregation levels encode proteins that act as positive regulators of transcription (Fig. S1; Table S1) raising the possibility that a decrease in global transcription, and hence protein synthesis, limits protein aggregation levels during As(III) stress. To test this, we incubated cells with As(III) and the transcription inhibitor 1,10-phenanthroline, added either separately or together, and monitored the cellular distribution of Hsp104-GFP. Indeed, 1,10-phenanthroline strongly attenuated As(III)-induced protein aggregation as few Hsp104-GFP foci were detected in the presence of this chemical (Fig. 5A). Next, we scored protein aggregation levels in cells lacking *RPB4* encoding a subunit of the RNA polymerase II enzyme; this mutant has been shown to have a decreased global transcription while it maintains a balanced level of mRNA because it is globally stabilized [51]. The *rpb4Δ* mutant was transformed with Sis1-GFP, an essential Hsp70 co-chaperone [52] that associates with aggregation-prone proteins under conditions of proteotoxic stress [53–55]. As for Hsp104-GFP, 1 h of exposure to As(III) resulted in Sis1-GFP redistribution to distinct cytosolic foci, with the majority of cells containing several foci (Fig. 5B). About 80% of wild type cells contained Sis1-GFP foci after 1 h of exposure but the total fraction of cells with foci decreased over time, as did the proportion of cells containing ≥3 Sis1-GFP foci/cell. Interestingly, *rpb4Δ* had substantially less aggregates than the wild type during exposure (Fig. 5B). Thus, a global reduction or inhibition of transcription by chemical or genetic means can mitigate protein aggregation during As(III) stress. To test whether a global reduction of transcription is accompanied by reduced translation, we performed polysome profiling assays in wild type and *rpb4Δ* cells that were either untreated or exposed to As(III). For wild type cells, a general repression of translation initiation occurs in response to As(III) evidenced by increased levels of ribosomal subunits (40S and 60S) and monosomes (80S), and by decreased levels of polysomes (Fig. 5C). Quantification of the polysome-monosome ratio (P/M) in wild type cells showed a ~2-fold reduction in translation during exposure. Importantly, the *rpb4Δ* mutant had lower translational activity than the wild type both in the absence and presence of As(III) (Figs. 5C). The P/M ratio for *rpb4Δ* was about 2-fold lower than for wild type cells in the absence of stress and was further reduced by ~30% during As(III) exposure. Importantly, *rpb4Δ* cells grew better in As(III)-containing medium than wild type cells (Fig. 5D) indicating that reduced protein synthesis is beneficial during As(III) exposure. In contrast, the *rpb4Δ* mutant was more sensitive to increased temperature (37°C), a condition that causes misfolding of nascent as well as native proteins. The *rpb4Δ* mutant was also sensitive to high osmolarity (1M NaCl), a condition that does not affect protein folding. Taken together, for *rpb4Δ* a reduction in global transcription is accompanied by low translational activity that likely protects the *rpb4Δ* proteome from As(III)-induced misfolding and aggregation.

**Figure 5.**
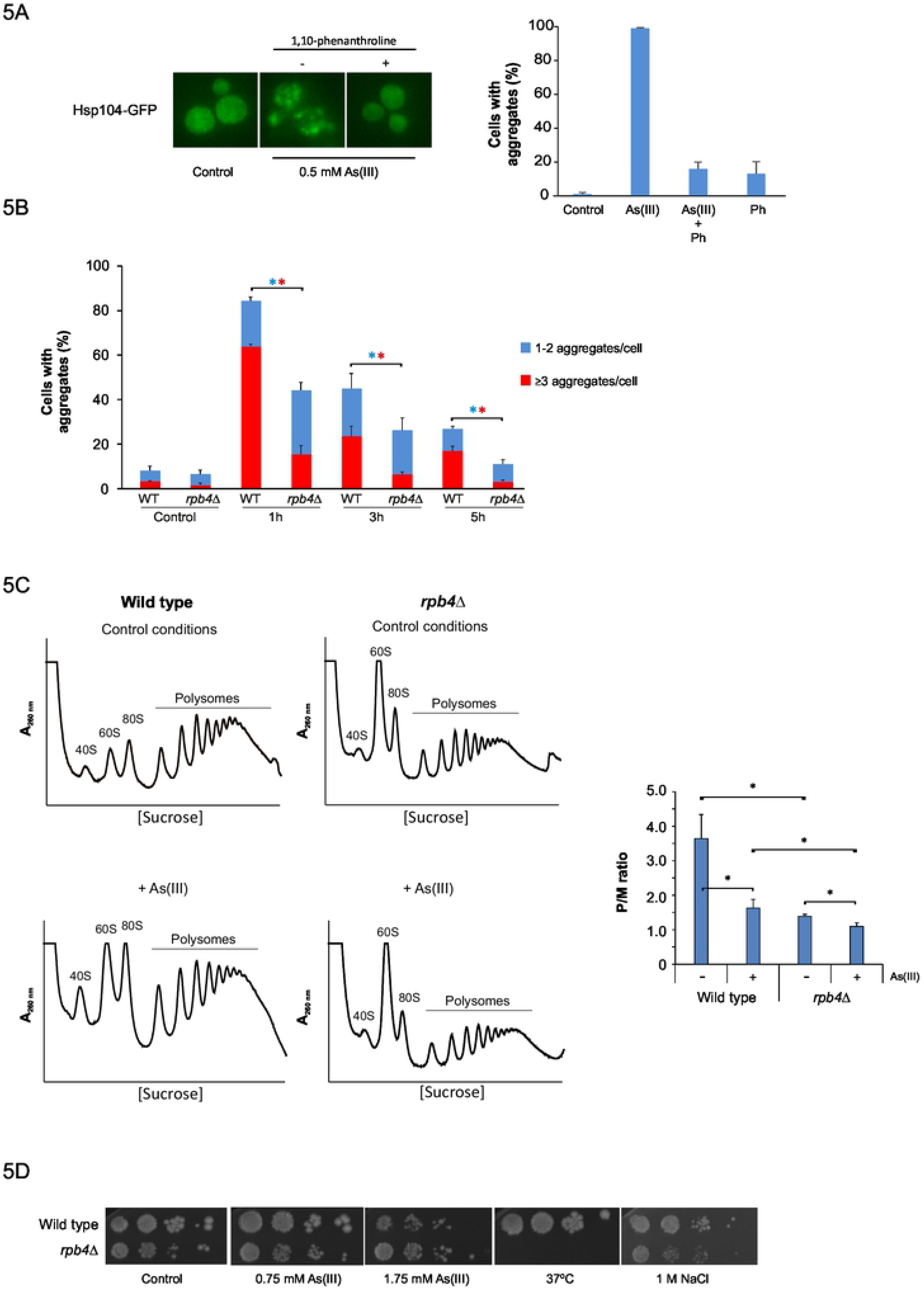
Global transcription affects protein aggregation levels during As(III) stress. **5A)** Inhibition of transcription attenuates As(III)-induced aggregate formation. Hsp104–GFP localization was monitored in wild type cells before and after 1 h exposure to 0.5 mM As(III) in the absence or presence of 0.1 mg/ml 1,10-phenantroline (Ph). The fractions of cells containing aggregates were determined by visual inspection of 20-135 cells per condition. Error bars represent S.D. from three independent biological replicates (n=3). Left panel: representative images are shown. Right panel: quantification of protein aggregation. **5B)** Quantification of protein aggregation. Sis1-GFP distribution was scored in wild type and *rpb4Δ* cells by fluorescence microscopy before and after exposure to 0.5 mM As(III). The fractions of cells containing 1-2 aggregates/cell and ≥3aggregates/cell were determined by visual inspection of 246-370 cells per condition and time-point. Error bars represent standard deviations (S.D.) from three independent biological replicates (n=3). The error bars on the top concern the total fraction of cells with aggregates, whilst those on the red bars concern the fraction of cells with ≥3 aggregates/cell. Statistical analyses were performed by Student’s t-test and significant differences are indicated by star (*p*<0.05). Blue star: total fraction of cells with aggregates; red star: ≥3 aggregates/cell. **5C)** Translational response to As(III) exposure. Polysome profiles were obtained from wild type and *rpb4Δ* strains cultivated in synthetic complete medium for at least 3 h to reach exponential phase, and then cells were further maintained in or treated with 0.75 mM As(III) for 1 h. The A_260nm_ profiles after gradient fractionation are shown and the ribosomal subunits (40S and 60S), monosomes (80S) and polysomes are indicated. Three biologically independent replicates were performed in each case (n=3). A representative profile is shown. Right panel: the average polysome/monosome ratio (P/M) is represented with its corresponding standard deviation. Statistical analyses were performed by Student’s t-test and star indicates statistically significant difference (*p*<0.05). **5D)** 10-fold serial dilutions of the indicated cells were plated onto YPD agar plates with or without As(III) or NaCl. Growth was monitored after 2-3 days at 30°C or 37°C.

### Loss of transcriptional control impacts protein aggregation levels during As(III) stress through distinct mechanisms

It is unlikely that all transcription-related hits identified in this current study affect global transcription as shown for Rpb4. Instead, the majority of these proteins are expected to have limited impact on global mRNA levels and likely affect proteostasis through distinct mechanisms.

#### Positive regulators of transcription

We asked whether transcription-related mutants with reduced aggregation levels identified in our screen exhibit a general resistance to proteotoxic stress. Therefore, we tested growth of mutants lacking selected positive regulators of transcription (Msn4, Spt3, Srb2, Met18, Gat4, Ace2, Spt8) in the presence of hygromycin B (HygB), a chemical that reduces translational fidelity leading to increased misincorporation of amino acids into nascent polypeptides causing them to misfold [56]. The activator Msn4 identified in our screen has a paralogue Msn2; Msn2 and Msn4 are partially redundant and they induce gene expression in response to several types of stress, including As(III) [57–59]. We therefore included the *msn2Δ* mutant as well as the *msn2Δ msn4Δ* double mutant. Growth assays showed that *met18Δ*, *ace2Δ* and *msn2Δ msn4Δ* cells were HygB resistant (Fig. 6A). Hence, deletion of these positive regulators of transcription might mitigate proteotoxicity. Western blot analyses showed that all mutants had similar levels of stress-induced Hsp104 and HSP70 as wild type cells after As(III) treatment (Fig. S3), suggesting that the reduced aggregation levels during As(III) stress observed in these mutants is not a result of high levels of molecular chaperones.

Msn2 and Msn4 were previously shown to be activated and to control induced expression of ~60 genes during As(II) stress [59]. Therefore, the anticipated effect of *MSN2/MSN4* deletion would be a decreased ability to cope with As(III) stress due to a compromised stress response. Instead, deletion of both genes results in increased tolerance to As(III) (Fig. 6B) [59], whereas overexpression of either *MSN2* or *MSN4* from a strong constitutive promoter enhances sensitivity [59]. To substantiate our data that suggest diminished proteotoxic stress in *msn2Δ msn4Δ*, we scored protein aggregation levels in *msn2Δ msn4Δ* cells carrying Sis1-GFP. Importantly, after 3 and 5 h of exposure the total fraction of cells with aggregates as well as the fraction of cells with ≥3 Sis1-GFP foci/cell was clearly lower in *msn2Δ msn4Δ* compared to the wild type (Fig. 6C). In a reciprocal assay, moderate overexpression of *MSN2* or *MSN4* (native promoters; centromeric plasmids) in wild type cells carrying Hsp104-GFP resulted in a higher fraction of cells with aggregates/Hsp104-GFP foci than cells transformed with the empty vector after 1 h of As(III) exposure (Fig. 6D). A higher fraction of cells overexpressing *MSN4* contained aggregates also at the 3 h time-point compared to cells transformed with the empty vector. Hence, *MSN2/4* gene dosage has a substantial impact on proteostasis. Since As(III) targets nascent proteins for aggregation [18], we examined translational activity in *msn2Δ msn4Δ* cells by polysome profiling. In the absence of As(III), the translational activity was similar for wild type and *msn2Δ msn4Δ* cells (Fig. 6E). Interestingly, lack of the stress-responsive activators Msn2 and Msn4 resulted in a strong inhibition of translation initiation in response to As(III). Quantifying the polysome-monosome ratio (P/M) showed that *msn2Δ msn4Δ* cells were about 2-fold more efficient in reducing translation during As(III) exposure than wild type cells (Fig. 6E). Thus, the ability to efficiently decrease translation is likely responsible for the diminished protein aggregation levels observed in *msn2Δ msn4Δ* cells, as well as for its HygB and As(III) resistance.

#### Negative regulators of transcription

We next turned to a subset of transcription-related hits with enhanced aggregation levels encoding proteins that act as transcriptional repressors (Mig1, Ure2), are components of the Rpd3L histone deacetylase complex (Dep1, Rxt2, Rxt3, Sap30) or are involved in chromatin silencing (Esc2, Rlf2, Hst3) (Fig. S2; Table S1). To test whether mutants with enhanced aggregation levels exhibit sensitivity during proteotoxic stress, we scored growth of these mutants in the presence of HygB. We also included a mutant lacking the ribosome-associated chaperones Ssb1 and Ssb2 (*ssb1/2Δ*) as a positive control [56]. All tested mutants (*dep1Δ*, *esc2Δ*, *rlf2Δ*, *rxt2Δ* and *mig1Δ*) were sensitive to HygB (Fig. 7A), suggesting that they might experience enhanced proteotoxic stress. To substantiate this, we took a genetic approach by introducing *SSE1* deletion into the *dep1Δ*, *esc2Δ*, *rlf2Δ*, *rxt2Δ* and *mig1Δ* cells and testing whether this molecular chaperone is critical for their growth. Sse1 acts as a nucleotide exchange factor for HSP70 chaperones [60, 61], is required for Hsp104-dependent protein disaggregation [62], and cells lacking Sse1 have been shown to have enhanced protein aggregation levels [63]. There was a clear As(III)-dependent synthetic growth defect in all tested double mutants (Fig. 7B). In addition, the double *dep1Δ sse1Δ* and *rxt2Δ sse1Δ* mutants grew poorly already in the absence of As(III) and this growth defect was strongly aggravated in the presence of metalloid. These observations suggest that lack of these negative regulators of transcription sensitizes cells to protein folding stress. We confirmed enhanced protein aggregation levels in cells lacking *MIG1*, encoding a protein involved in glucose repression [64]; the total fraction of cells with aggregates as well as the fraction of cells with ≥3 Sis1-GFP foci/cell was significantly higher in *mig1Δ* than the wild type at all time-points during exposure (Fig. 7C). Western blot analyses showed that all mutants had lower levels of Hsp104 compared to wild type cells during As(III) exposure although some mutants (*esc2Δ*, *rfl2Δ*) had slightly elevated Hsp104 levels in the absence of stress (Fig. S4). HSP70 levels were similar in wild type and the mutants (Fig. S4). Hence, an inability to properly adjust chaperone levels might underlie the increase in protein aggregation observed in these mutants. Collectively, the results above indicated that loss of transcriptional control impacts PQC through distinct mechanisms including translational control and protein folding.

### Translational repression is central for proteostasis and cell viability during As(III) stress

The results above indicated that ongoing protein synthesis is detrimental during As(III) exposure. In particular, reduced translation was associated with less aggregates and improved growth for the *rpb4Δ* and *msn2Δ msn4Δ* mutants (Figs. 5,6). Thus, translational repression might be important to control proteostasis during As(III) stress. To substantiate this, we measured protein aggregation levels in a selection of mutants lacking proteins with functions in translational repression and mRNA decay including Dhh1, Xrn1, Not1 and Ccr4 [65–67]. Indeed, all tested mutants had elevated protein aggregation levels during As(III) exposure; the total fraction of cells with Sis1-GFP foci/aggregates as well as the fraction of cells with ≥3 Sis1-GFP foci/cell was significantly higher in *dhh1Δ, xrn1Δ*, *ccr4Δ* and *not1Δ* cells compared to wild type at the 3 and 5 h time-points (Fig. 8A). In addition, *xrn1Δ*, *ccr4Δ* and *not1Δ* cells had higher levels of aggregates also in the absence of stress. Cells lacking *DHH1* have been shown to be defective in translational repression during glucose starvation [65]. Likewise, polysome profiling during As(III) exposure showed that *dhh1Δ* cells are ~1.5-fold less efficient than wild type cells in inhibiting translation (Fig. 8B). Growth assays indicated that cells lacking *DHH1*, *XRN1*, *CCR4* and *NOT1* are As(III) sensitive (Fig. 8C). Thus, the inability to efficiently repress translation is likely responsible for the enhanced protein aggregation levels and As(III) sensitivity observed in these mutants.

**Figure 8.**
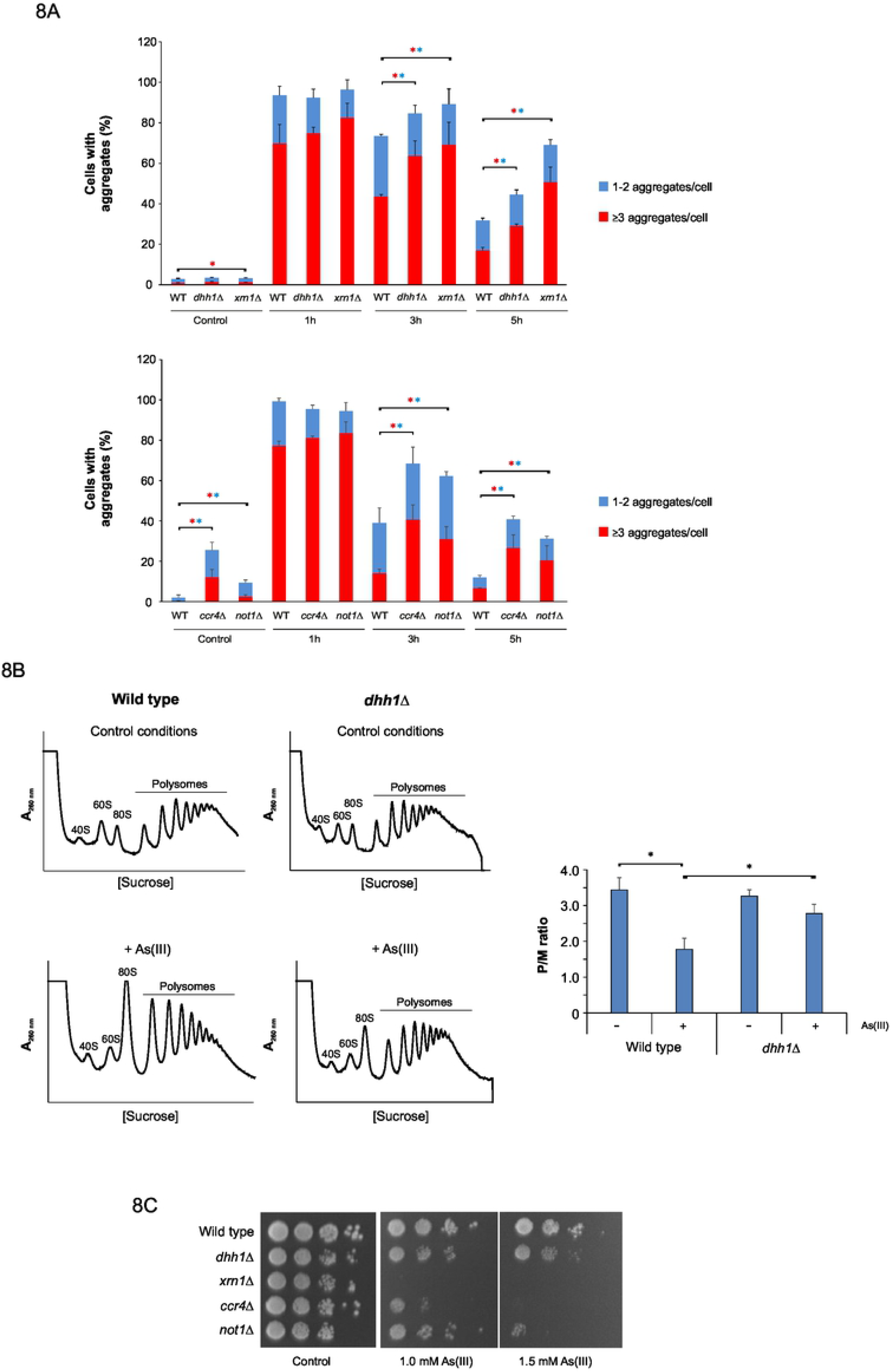
Translational repression is central for proteostasis and cell viability during As(III) stress. **8A)** Quantification of protein aggregation. Sis1-GFP distribution was scored in wild type and mutants defective in translational repression by fluorescence microscopy before and after exposure to 0.5 mM As(III). The fractions of cells containing 1-2 aggregates/cell and ≥3aggregates/cell were determined by visual inspection of 121-361 cells per condition and time-point. Error bars represent standard deviations (S.D.) from three biological replicates (n=3). The error bars on the top concern the total fraction of cells with aggregates, whilst those on the red bars concern the fraction of cells with ≥3 aggregates/cell. Statistical analyses were performed by Student’s t-test and significant differences are indicated by star (*p*<0.05). Blue star: total fraction of cells with aggregates; red star: ≥3 aggregates/cell. **8B)** Cells lacking Dhh1 are defective in translational repression during As(III) stress. Polysome profiles were obtained from wild type and *dhh1Δ* strains cultivated in synthetic complete medium for at least 3 h to reach exponential phase, and then cells were further maintained in or treated with 0.75 mM As(III) for 1 h. The A_260nm_ profiles after gradient fractionation are shown and the ribosomal subunits (40S and 60S), monosomes (80S) and polysomes are indicated. Three biologically independent replicates were performed in each case (n=3). A representative profile is shown. Right panel: the average polysome/monosome ratio (P/M) is represented with its corresponding standard deviation. Star indicates statistically significant difference (*p*<0.05). **8C)** 10-fold serial dilutions of the indicated strains were plated onto YPD agar plates with or without As(III). Growth was monitored after 2-3 days at 30°C.

## DISCUSSION

### A comprehensive view of PQC during arsenite stress

Arsenic is a highly poisonous and carcinogenic metalloid that causes widespread protein misfolding and aggregation [17, 18, 23]. Elucidating how protein aggregates are formed *in vivo*, the mechanisms by which they affect cells, and how cells prevent their accumulation is important for understanding arsenic’s toxicity and pathogenicity. Our genome-wide imaging screen uncovered novel genes and processes that are crucial for proteostasis during As(III) stress (Table S1). Mutants with reduced aggregation levels are enriched for functions related to protein biosynthesis and transcription, whilst functions related to cellular signalling, metabolism, and protein folding and degradation are overrepresented among mutants with enhanced aggregation levels (Fig. 2; Table S1). Several of the systems identified were known or anticipated to influence PQC, validating the screening approach. For example, a set of hits with enhanced levels of protein aggregation are associated with protein folding, turn-over and degradation. Thus, compromising the cellular capacity to correctly fold or degrade proteins increases the burden of misfolded and aggregated proteins during As(III) stress. We also identified a large number of genes and networks which were previously not linked to proteostasis, including proteins with functions related to signalling, chromatin organization and transcription. The mechanistic details by which many of the identified genes contribute to PQC remain to be established. Here, we have chosen to focus on selected hits from the screen and highlighted the importance of transcriptional and translational control as well as of restricting intracellular arsenic to mitigate proteotoxicity.

### Protein aggregation is correlated with intracellular arsenic levels and As(III) toxicity

Genes related to arsenic transport and detoxification were present in both the enhanced and reduced aggregation sets. Loss of proteins that restrict cytosolic arsenic levels resulted in enhanced protein aggregation, whilst loss of proteins that contribute to arsenic influx lead to less aggregates. For example, cells lacking the As(III) entry pathway Fps1 or its regulators Rgc1 and Rgc2 accumulated less intracellular arsenic and showed reduced aggregation levels (Fig. 3). The *fps1Δ* and *rgc1Δ rgc2Δ* mutants are As(III) resistant [39, 40]. Conversely, the absence of the As(III) exporter Acr3 or its transcriptional regulator Yap8 caused As(III) sensitivity, increased levels of intracellular arsenic and enhanced levels of protein aggregation [42, 44] (Fig. 3). Likewise, deletion of *YCF1*, important for vacuolar sequestration of arsenic, or its transcriptional regulator *YAP1* lead to increased levels of intracellular arsenic, protein aggregation, and As(III) sensitivity [42, 44] (Fig. 3). Our data implicated the Hog1 MAP kinase pathway in PQC. Hog1 is a negative regulator of Fps1 and deletion of *HOG1* results in increased As(III) influx [47], enhanced protein aggregation (Fig. 3), and As(III) sensitivity [47]. Deleting negative regulators of Hog1 (Ptp2 and Ptp3) resulted in less intracellular arsenic, less aggregates (Fig. 3) and enhanced resistance [47]. Thus, Hog1 and HOG pathway components appear to contribute to PQC primarily by modulating Fps1 activity and intracellular arsenic levels. Collectively, these data strongly suggest that intracellular arsenic is a direct cause of protein aggregation and that restricting the cellular accumulation of this metalloid is an important means for cells to protect their proteome from As(III)-induced damage and toxicity. Indeed, cells respond to As(III) by downregulating arsenic influx pathways [39, 68] and by increasing expression of arsenic export and sequestration systems [48, 69]. An important aspect of future work will be to systematically measure intracellular arsenic in all ~400 mutants identified in this study to pin-point pathways and regulators associated with arsenic uptake, efflux, intracellular distribution and sequestration.

Previous data suggested that protein aggregation may contribute to As(III) toxicity. This was based on, amongst others, the correlation of protein aggregation levels and As(III) sensitivity/resistance in a limited set of mutants [18]. Our current study provides genome-wide support for this notion. We found a strong correlation between mutants that were As(III) sensitive and mutants with enhanced protein aggregation during exposure. Likewise, there was a strong correlation between mutants that were As(III) resistant and those that showed less protein aggregation (Fig. 2). Previous studies indicated that proteins are the main target of As(III) during acute exposure [48] and that hundreds of proteins aggregate [18, 23]. Misfolded forms of these proteins might engage in extensive aberrant protein-protein interactions, resulting in an increased burden on PQC systems and an impact on cell viability [18, 23]. Together, these findings support the notion that protein aggregation contributes to As(III) toxicity and that aggregate management is crucial for cell survival.

### Appropriate transcriptional control is important for proteostasis and As(III) resistance

The importance of transcriptional control for proteostasis has been highlighted only in a few studies. For example, transcription errors have been shown to induce proteotoxic stress and to shorten lifespan in yeast [49]. Here, we show that As(III) does not cause errors at the level of transcription (Fig. 4). As(III)-exposed cells downregulate gene expression associated with protein synthesis and upregulate expression of genes related to protein folding and degradation [23, 69, 70]. Moreover, yeast cells decrease the expression of aggregation-prone proteins during As(III) exposure [23], possibly as a means to mitigate aggregation and toxicity. Here, we provide additional evidence that appropriate transcriptional control is crucial for PQC during As(III) exposure. First, a large number of transcription-related hits were found in both the enhanced and reduced aggregation sets. These hits encode positive as well as negative regulators of transcription (Table S1, Fig. 2). Selected mutants lacking transcriptional repressors accumulated more aggregates than wild type cells and were As(III) sensitive. Genetic and biochemical data indicated that some of these mutants experience proteotoxic stress in the absence of As(III), and that protein aggregation is exacerbated during exposure. The molecular chaperone Sse1 was required to sustain growth of these mutants, supporting the notion that they suffer from proteotoxic stress (Fig. 7). Similarly, selected mutants lacking transcriptional activators showed resistance to HygB, suggesting that their absence might mitigate proteotoxicity (Fig. 6). Second, our data suggest that global transcription affects protein aggregation levels during As(III) stress. Inhibition of transcription with 1,10-phenantroline attenuated As(III)-induced aggregate formation (Fig. 5), which is similar to the effect observed with cycloheximide, a chemical inhibitor of translation [18]. Moreover, protein aggregation levels were reduced in the *rpb4Δ* mutant that has decreased global transcription rate [51]. Thus, transcriptional inhibition by genetic or chemical means mitigates As(III)-induced protein aggregation. For some of the identified mutants, it is possible that altered transcription also affects global translation, *i.e.* by altering the number of nascent proteins synthesized that can be targeted by As(III) to misfold and aggregate. This was indeed the case for *rpb4Δ*; polysome profiling assays demonstrated that *rpb4Δ* had lower translational activity than the wild type both in the absence and presence of As(III) (Fig. 5). However, most of the transcription-related mutants identified are likely to affect aggregation through other mechanisms, since several of these hits regulate expression of a limited set of target genes and are therefore expected to have little impact on global mRNA levels. Instead, the identified regulators might affect a variety of gene targets that influence the cellular capacity to either deal with the amounts of intracellular As(III) and/or the damages caused by the metalloid. We showed that some of the transcription-related mutants tested had lower levels of molecular chaperones (*dep1Δ*, *esc2Δ*, *rlf2Δ*, *rxt2Δ*, *mig1Δ*), accumulated high levels of intracellular arsenic (*yap1Δ*, *yap8Δ*) or showed altered capacity to control translation (*msn2Δ msn4Δ*). Interestingly, the mutant lacking the stress-activated factors Msn2 and Msn4 was resistant to As(III) and HygB, and displayed less protein aggregation during As(III) stress compared to wild type cells (Fig. 6). This finding was unexpected since Msn2/Msn4 regulate As(III)-induced expression of ~60 genes [59]. It was previously hypothesized that a chronic activation of general stress factors by Msn2/Msn4 may contribute to As(III) sensitivity [59]. Here, we demonstrate that lack of Msn2 and Msn4 promotes strong translation inhibition in response to As(III) (Fig. 6). Hence, an improved efficiency of translation inhibition is likely responsible for the diminished aggregation levels observed in *msn2Δ msn4Δ*, as well as for its HygB and As(III) resistance. Collectively, these data support the notion that accurate transcriptional control is crucial to protect cells from the accumulation of misfolded and aggregated proteins. The mechanisms by which transcriptional regulators impact PQC appear to be distinct.

### Translational control is crucial to mitigate As(III)-induced proteotoxicity

Our data highlights that translational control is crucial for aggregate management and As(III) resistance. First, a large proportion of the mutants with reduced aggregation (approximately 25% of the hits) are part of highly connected networks involved in cytoplasmic translation (*e.g.* ribosomal proteins), rRNA modification and processing, and ribosome assembly (Figs. 2,S1). These mutants may have decreased translational activity, suggesting that ongoing translation results in As(III)-induced protein aggregation. This finding is in line with previous studies showing that proteins are particularly susceptible to As(III)-induced aggregation during translation/folding [18, 23] and that proteins with high translation rates are particularly susceptible for aggregation during As(III) stress [23, 71]. Second, we demonstrate that diminished translation mitigates As(III)-induced protein aggregation and toxicity. Polysome profiling assays showed that wild type cells repress translation during As(III) exposure (Fig. 5). This is likely a result of increased eIF2α phosphorylation [31], inhibition of translation initiation [72] and reduction of ribosomal protein levels [31]. Importantly, repression of translation appears crucial for safeguarding the proteome from As(III)-induced damage. Indeed, low translational activity of *rpb4Δ* protected this mutant from As(III)-induced protein aggregation and toxicity (Fig. 5). Similarly, an improved efficiency in translational repression, as shown for *msn2Δ msn4Δ* cells, mitigated protein aggregation and resulted in As(III) resistance (Fig. 6). Our data also emphasized the key role of translation initiation control in response to As(III) in contrast to other proteotoxic stresses like heat shock (Fig. 5E). Third, we demonstrate that mutants lacking proteins with functions in translational repression have enhanced protein aggregation levels and exhibit As(III) sensitivity (Fig. 8). Polysome profiling assays demonstrated that *dhh1Δ* was indeed defective in translational repression during As(III) stress. Thus, low translation and efficient translation repression is beneficial during As(III) exposure whilst lack of translational repression is detrimental for the cell. Together, these data strongly support the notion that appropriate control at the translational level is crucial to mitigate As(III)-induced proteotoxicity.

### Implications for human disease processes

Previously, we found several homologues of yeast proteins that aggregated during As(III) exposure to be present in human disease-associated aggregates in AD, PD and amyotrophic lateral sclerosis (ALS), suggesting that the mechanisms underlying protein aggregation in stress-exposed yeast cells may be relevant during human disease processes [23, 71]. The hit list from the current screen is also interesting in the light of human pathogenesis as several homologues to proteins related to human diseases were identified (Table S1). For example, lack of Tdp1, encoding tyrosyl-DNA phosphodiesterase I, resulted in increased protein aggregation levels. Mutation in the human homologue, TDP1, results in the inherited neurodegenerative disorder SCAN1 [73]. Thus, SCAN1 patients may be susceptible for conditions of proteotoxic stress, such as chronic exposure to low levels of metals. Similarly, lack of *LTN1* resulted in elevated aggregation levels during As(III) exposure. Ltn1 is the yeast homologue of mammalian Listerin, an E3 ubiquitin ligase implicated in neurodegeneration [74]. The screen also identified several components of the elongator complex (ELP1-6) that is primarily involved in tRNA modifications [28, 75]; deletion of *ELP1*/*IKI3*, *ELP2*, *3*, *4* or *6* caused reduced levels of protein aggregates during As(II) exposure. Mutations in the human Elp1 homolog IKAP causes familial dysautonomia, a rare neurodegenerative disease that is associated with growth abnormalities and degradation of sensory functions, and patients suffering from this disease showed reduced levels of wobble uridine tRNA modification [75]. Elp3 is suggested to be a modifier of ALS, a fatal degenerative motor neuron disorder, indicating a possible link between tRNA modifications and neurodegeneration [76]. Thus, members of the elongator complex are implicated in disease processes as well as PQC during metalloid exposure. There is evidence that chronic heavy metal and metalloid exposure may promote the progression of certain neurodegenerative and age-related disorders, such as AD and PD [12–15]; however, the underlying mechanisms remain largely unknown. A better understanding of the role of the proteins above in PQC may shed novel light on disease processes associated with metal exposure.

## Conclusions

Collectively, this work provides a comprehensive view of the cellular machineries involved in As(III)-dependent protein aggregate management and highlights the importance of transcriptional and translational control. The broad network of cellular systems that impinge on proteostasis during As(III) stress identified in this current study provides a valuable resource and a framework for dedicated follow-up studies of the molecular underpinnings of arsenic toxicity and pathogenesis.

## MATERIALS AND METHODS

### Yeast strains and culturing conditions

A list of *S. cerevisiae* strains used in this study is presented in Table S2. Most strains are based on the BY4741 and BY4742 laboratory strains [77], W303-1A [78] and the yeast deletion collection [79]. Double mutants were generated by crossing haploid single mutants using standard procedures and all double mutants were confirmed by PCR. Yeast strains were routinely grown on minimal SC (synthetic complete) medium (0.67% yeast nitrogen base) supplemented with auxotrophic requirements and 2% glucose as a carbon source or in rich YPD (yeast peptone dextrose) medium. Growth assays were carried out on solid agar as previously described [44]. Sodium arsenite (NaAsO_2_), 1,10-phenantroline (both from Sigma-Aldrich) and hygromycin B (Formedium) were added to the cultures at the indicated concentrations.

### Yeast library creation and automated high-content microscopy

The yeast library harbouring the Hsp104-GFP aggregate marker was created as previously described [80] by crossing the Hsp104-GFP query strain into a genome-wide collection of viable yeast single deletion mutants (SGA-v2, Boone lab) using a synthetic genetic array (SGA) approach [81, 82]. Cells from the collection were transferred into 96-well plates using a liquid handling robot (Microlab STAR, Hamilton Company), and treated with 0.25 mM As(III) for 2 h to induce protein aggregation, followed by fixation with 3.7% formaldehyde. Then cells were washed 2 times in 1xPBS (phosphate-buffered saline) and transferred into 96-well glass bottom plates for imaging. Automatic image acquisition of treated cells was performed using a high-content microscope (ImageXpress MICRO, Molecular Devices). 25 images were acquired for each mutant. Quantification of number of cells with aggregates and number of aggregates per cell was performed using MetaXpress (version 3, Molecular Devices) software.

### Bioinformatics and network analyses

Categories of over-represented protein functions in our data-sets were identified using FunCat at Munich Information Center for Protein Sequences (MIPS) [25]. Enriched functional categories were set with FDR < 0.05 using the Benjamini–Hochberg procedure [83]. Protein-protein interaction networks were constructed using the STRING database (string-db.org/) [26] with the organism set to *Saccharomyces cerevisiae* and the confidence score to the highest (0.9).

### Fluorescence microscopy

Yeast cells expressing the Hsp104-GFP (Table S2) or Sis1-GFP [53] fusion proteins were grown to mid-log phase in SC medium and treated with 0.5 mM As(III). Where indicated, 0.1 mg/ml 1,10-phenantroline was added. At the indicated time-points, cell samples were fixed with formaldehyde for 30 min at room temperature and washed with PBS. The GFP signals were observed using a Zeiss Axiovert 200M (Carl Zeiss MicroImaging) fluorescence microscope equipped with Plan-Apochromat 1.40 objectives and appropriate fluorescence light filter sets. Images were taken with a digital camera (AxioCamMR3) and processed with Zeiss Zen software. To quantify protein aggregation, the total fraction of cells with aggregates/Hsp104–GFP (or Sis1-GFP) foci as well as the fraction of cells having ≥3 aggregates/cell was determined using Image J software.

### Transcriptional errors

Transcription error assays were performed largely as described previously [50]. Patches of cells were grown on YPD medium overnight at 30°C, replica plated onto SC medium lacking histidine (His^−^) either with or without As(III) and incubated at 30°C for 3 days. Quantitative assay were performed in liquid medium. For this, cells were grown in 1 ml of YPD at 30°C overnight and then plated onto SC plates lacking histidine either with or without As(III) as well as on YPD plates to quantify the mean frequency of His^+^ cells relative to total number of viable cells. Plates were incubated at 30°C for 3 days and individual colonies were scored.

### Arsenic uptake

Intracellular arsenic was measured as described previously [47]. Briefly, exponentially growing cells were exposed to 1 mM As(III) for 1 h, collected and washed twice in ice-cold water. The cell pellet was then resuspended in water, boiled for 10 min, and centrifuged to collect the supernatant. The arsenic content of each sample was measured using a flame atomic absorption spectrometer (3300, Perkin Elmer) or by ICP-MS. Prior to analysis, the samples were diluted 20 times using water from a Thermo Scientific Barnstead GenPure water purification system (resistivity 18.2 MΩ cm) and acidified to 1% volume HNO_3_ (Sharlau, nitric acid 65% for trace analysis). The analysis was performed on an ICAP Q ICP-MS (Thermo Fisher Scientific) with an SC-FAST automated sampling introduction system (Elemental Scientific). The instrument was operated in the Kinetic Energy Discrimination (KED) mode with He as the collision gas to remove potential interference from argon chloride. Calibration was performed using a set of single element arsenic standards with concentrations up to 500 μg L^−1^. A 1 μg L^−1^ indium solution was continuously injected for internal standardization. The detection limit estimated based on 6 blank analyses was 0.1 μg As L^−1^.

### Protein analyses

Cells were grown in selective medium until log phase, after which cells were treated with 0.5 mM As(III). A volume of cells corresponding to OD_600_=1 was collected before and after 1 h of incubation with As(III), assuring the same amount of cells for each sample. Cells were treated with 2 M lithium acetate and incubated on ice for 5 min, centrifuged and the supernatant was discarded. A second treatment with 0.4 M NaOH was performed, followed by a 5 min incubation on ice. After centrifugation, cells were resuspended in 2X SDS loading buffer (125 mM Tris-HCl pH 6.7, 6% SDS, 2% Glycerol, 10% β-mercaptoethanol, bromophenol blue) and boiled for 5 minutes at 95°C. Protein samples were loaded on 10% TGX Stain-Free Gels and ran at 120V. Protein was transferred to a PVDF membrane using a Trans-Blot Turbo transfer system (Bio-Rad, Hercules, CA, USA), according to the manufacturer’s protocol. Membranes were blocked with 5% Blotting Grade Blocker (BioRad, Hercules, CA, USA)/Tris-buffered Saline – Tween 20 (5% BSA/TBS-T) (w/v), followed by an overnight incubation with the primary antibodies: anti-Hsp104 (anti-rabbit, ab69549, Abcam, Cambridge, U.K) or anti-Hsp70 (anti-mouse, ab47455, Abcam, Cambridge, U.K.). The following day, antibodies were washed three times with TBS-T, followed by a 1.5 h incubation with the respective secondary antibodies: StarBright700 anti-rabbit (10000068187, 1:5000) or StarBright700 anti-mouse (10000068185, 1:2500) both from BioRad (Hercules, CA, USA). Four washes with TBS-T were performed after incubation with secondary antibodies, after which the signal was detected using ChemiDoc (Bio-Rad, Hercules, CA, USA). The same protocol was performed for detection of the loading control, using the primary antibody anti-Pgk1 (anti-mouse, ab113687, Abcam, Cambridge, U.K.) and secondary antibody DyLight650 anti-mouse (84545, Invitrogen, Waltham, Massachusetts, USA). Images recovered from ChemiDoc were treated using the Image Lab Software (Bio-Rad, Hercules, CA, USA).

### Polysome profiling assays

To perform polyribosome profile analyses, a volume of an exponential growing cell cultures (OD_600_ between 0.6-1.0) corresponding to 50 OD_600_ units was collected. Then, cycloheximide (10 mg/ml) was added to a final concentration of 0.1 mg/ml and incubated 5 min on ice with occasional mixing. After centrifugation at 4.000 rpm for 5 min at 4°C, cells were resuspended and washed twice in 2 ml of cold lysis buffer (20 mM Tris-HCl pH 8.0, 140 mM KCl, 5 mM MgCl₂, 0.5 mM DTT, 0.1 mg/ml cycloheximide, 0.5 mg/ml heparin, 1 % Triton X-100). Then, cells were resuspended in 700 μl lysis buffer and transferred into a 2 ml screw-cap tube containing 500 μl of glass beads. Cells were broken by vortexing 8 times during 30 sec, with 30 sec of incubation on ice in between. After centrifugation (5.000 rpm, 5 min, 4°C), the supernatant was transferred into a new tube and centrifuged again (8.000 rpm, 5 min, 4°C). RNA from the supernatant was quantified. Glycerol was added to a final concentration of 5 % and samples were snap frozen and stored at -80°C. Samples were loaded onto 10–50% sucrose gradients and separated by ultracentrifugation for 2 h and 40 min at 35.000 rpm in a Beckman SW41Ti rotor at 4°C. The ultraviolet detection at A_260nm_ generated the general polyribosome profiles using Density Gradient Fractionation System and Isco UA-6 ultraviolet detector (Teledyne Isco, Lincoln, NE, USA).

## Acknowledgements

We thank Jeffrey Strathern (NCI Center for Cancer Research), Audrey Gasch (University of Wisconsin-Madison), Simon Alberti (Technical University Dresden), Elke Deuerling (University of Konstanz) and Stefan Hohman (Chalmers University of Technology) for providing strains and plasmids; Sebastien Rausch (Chalmers University of Technology) for the arsenic measurements; and José García-Martínez and José E. Pérez-Ortín (Universitat de València) for helpful suggestions to improve this work.

## Author contributions

Study conception and design: MJT, BL, TN; data collection and analysis: SA, AR, JIR, SH, XH, TJ, VK, AA, NB, MJT, BL; funding acquisition: MJT, TN, BL, AR. First draft was written by SA, AR, JIR, MJT. All authors commented on previous versions and read and approved the final manuscript.

## Supplemental figures and tables

**Fig S1:**
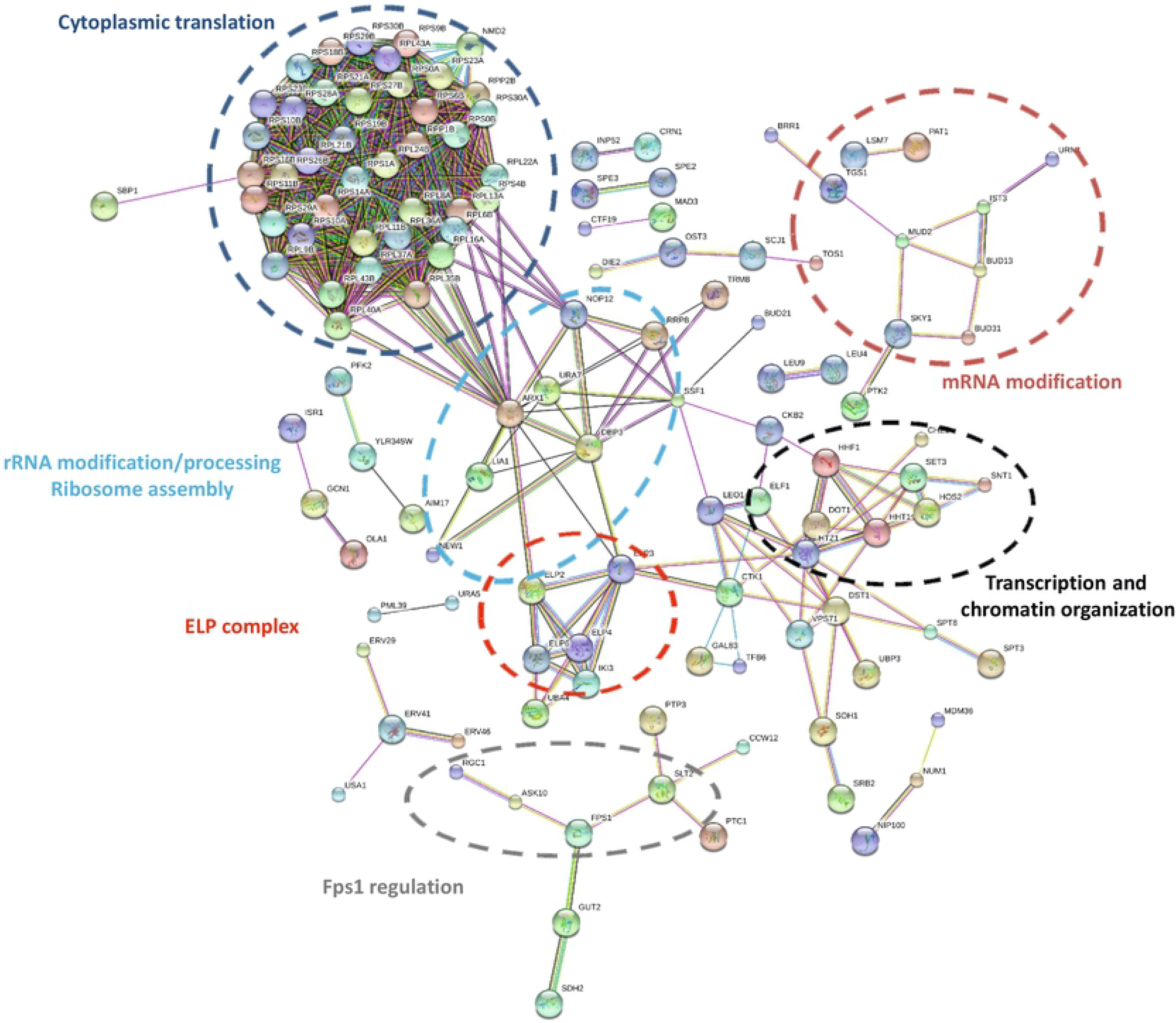
Protein-protein interaction networks among hits with reduced aggregation levels. Protein-protein interaction networks were constructed and visualized using the STRING database [26, 27] (evidence view, confidence score set to 0.9 (highest confidence). Only connected proteins are shown.

**Fig S2:**
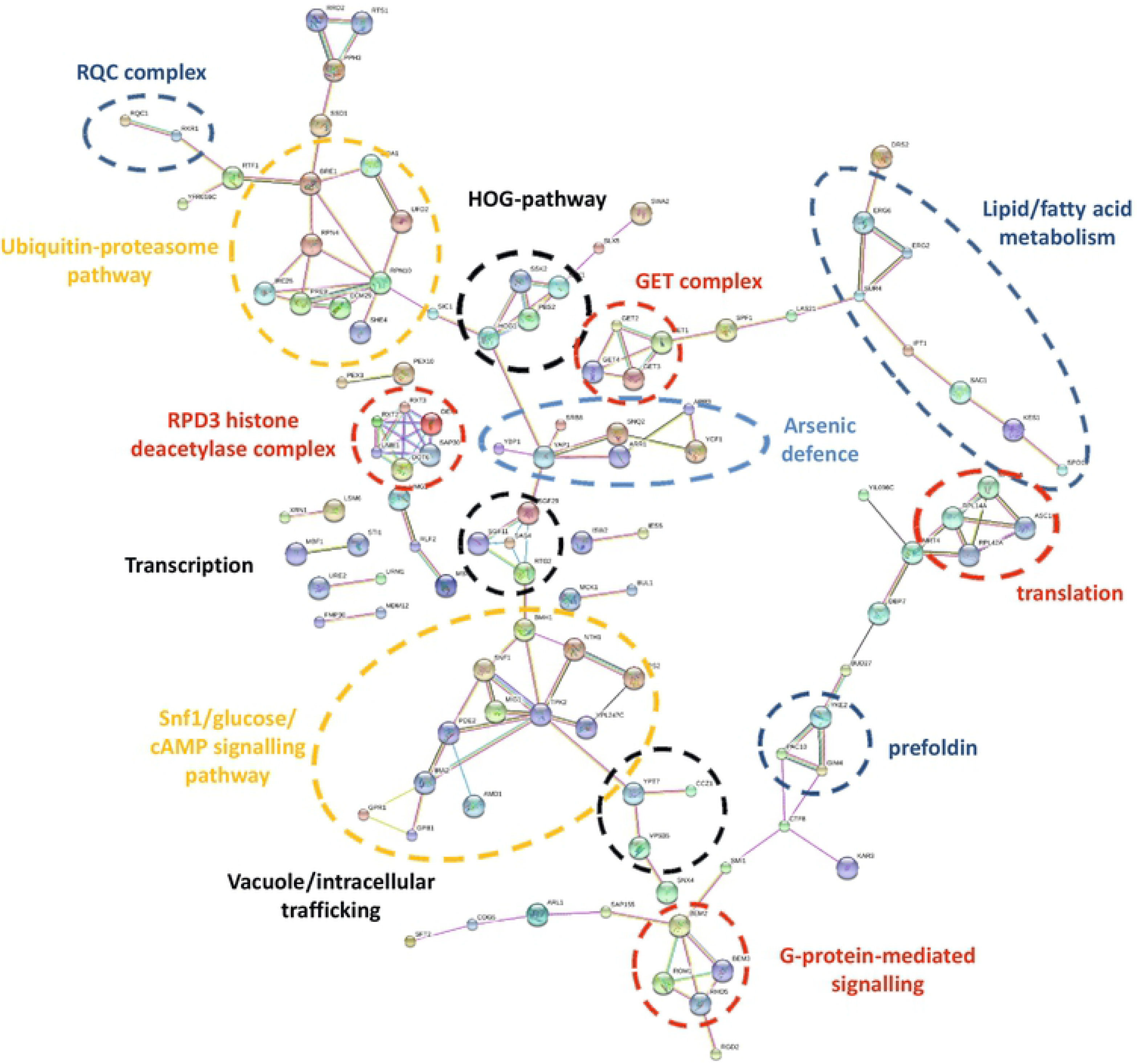
Protein-protein interaction networks among hits with enhanced aggregation levels. Protein-protein interaction networks were constructed and visualized using the STRING database [26, 27] (evidence view, confidence score set to 0.9 (highest confidence). Only connected proteins are shown.

**Fig S3:**
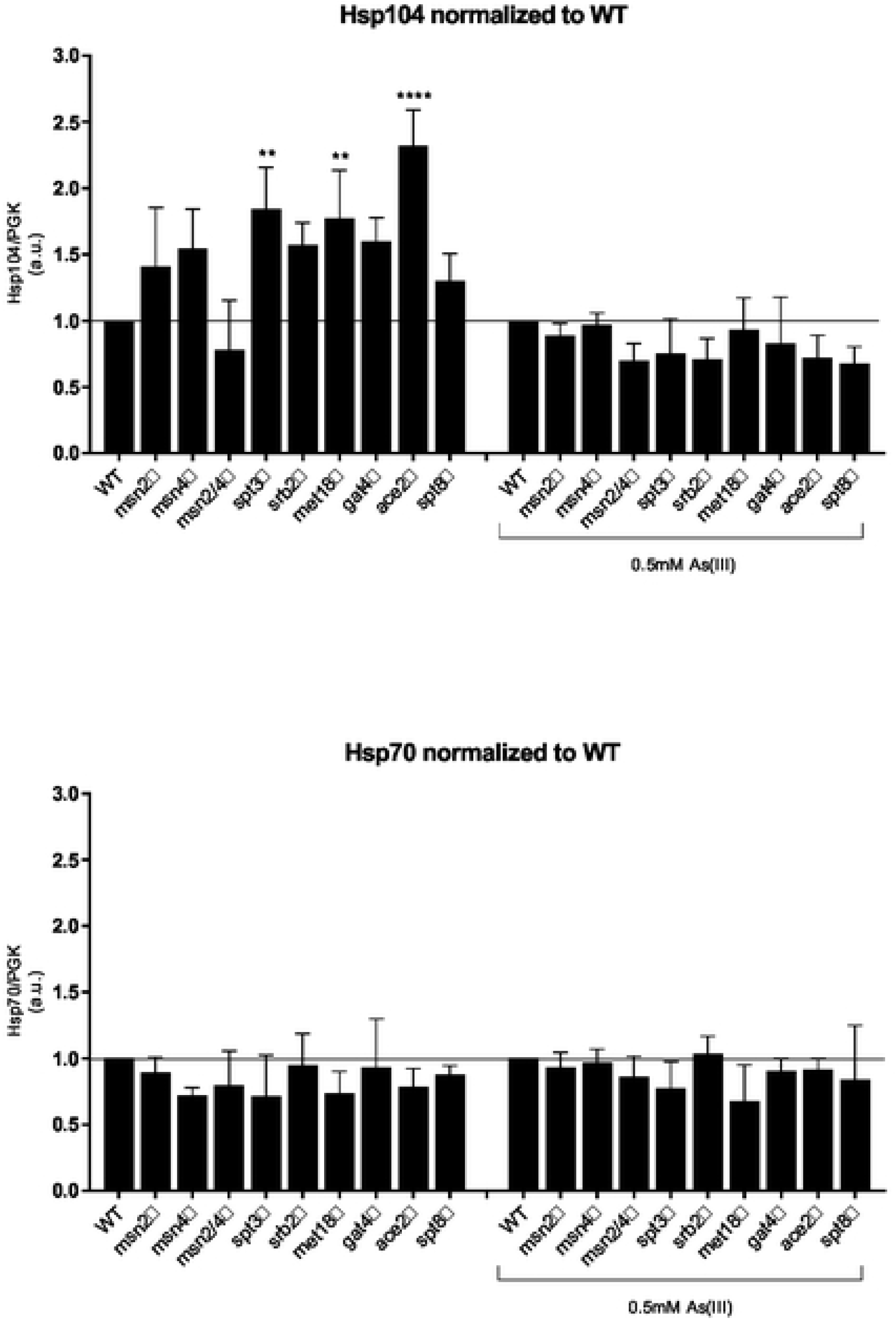
Hsp104 and Hsp70 protein levels in wild type cells and deletion mutants lacking positive regulators of transcription. Hsp104 and Hsp70 protein levels in wild type cells and deletion mutants lacking positive regulators of transcription before and after 1 h of 0.5 mM As(III) treatment. Hsp104 and Hsp70 levels were normalized to the levels of the loading control Pgk1. Protein levels were normalized to the wild type indicated by the horizontal line. Error bars represent standard deviations (S.D.) from four independent biological replicates (n=4). Statistical analyses were performed by One-Way ANOVA with Dunnett Test (\*\**p*<0.005, \*\*\*\**p*<0.00005)

**Fig S4:**
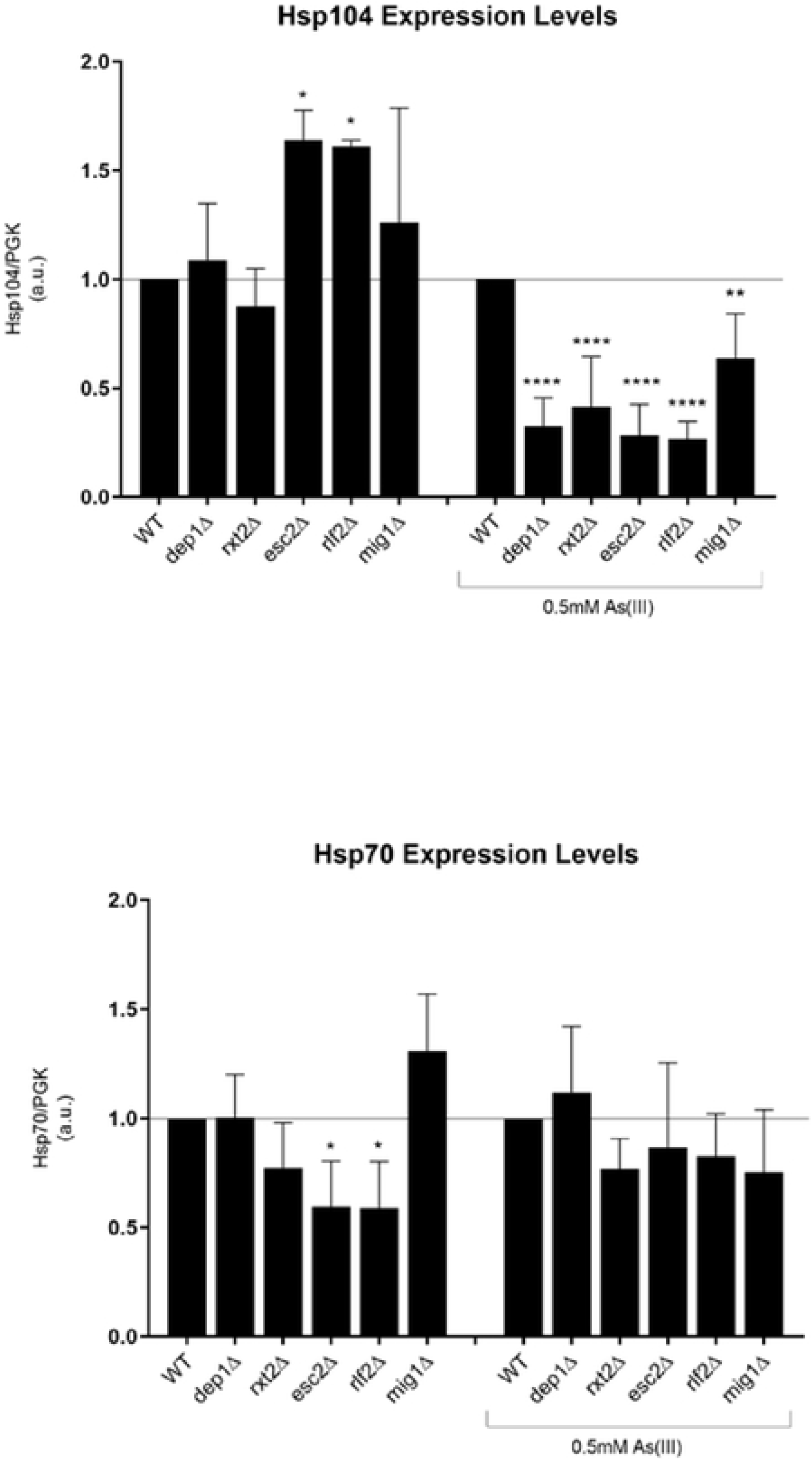
Hsp104 and Hsp70 protein levels in wild type cells and deletion mutants lacking negative regulators of transcription. Hsp104 and Hsp70 protein levels in wild type cells and deletion mutants lacking negative regulators of transcription before and after 1 h of 0.5 mM As(III) treatment. Hsp104 and Hsp70 levels were normalized to the levels of the loading control Pgk1. Protein levels were normalized to the wild type indicated by the horizontal line. Error bars represent standard deviations (S.D.) from six independent biological replicates (n=6). Statistical analyses were performed by One-Way ANOVA with Dunnett Test (\*\**p*<0.005, \*\*\*\**p*<0.00005)

